# Extensive mitochondrial population structure and haplotype-specific phenotypic variation in the *Drosophila* Genetic Reference Panel

**DOI:** 10.1101/466771

**Authors:** Roel P.J. Bevers, Maria Litovchenko, Adamandia Kapopoulou, Virginie S. Braman, Matthew R. Robinson, Johan Auwerx, Brian Hollis, Bart Deplancke

## Abstract

The *Drosophila* Genetic Reference Panel (DGRP) serves as a valuable resource to better understand the genetic landscapes underlying quantitative traits. However, such DGRP studies have so far only focused on nuclear genetic variants. To address this, we sequenced the mitochondrial genomes of >170 DGRP lines, identifying 229 variants including 21 indels and 7 frameshifts. We used our mitochondrial variation data to identify 12 genetically distinct mitochondrial haplotypes, thus revealing important population structure at the mitochondrial level. We further examined whether this population structure was reflected on the nuclear genome by screening for the presence of potential mito-nuclear genetic incompatibilities in the form of significant genotype ratio distortions (GRDs) between mitochondrial and nuclear variants. In total, we detected a remarkable 1,845 mito-nuclear GRDs, with the highest enrichment observed in a 40 kb region around the gene *Sex-lethal* (*Sxl*). Intriguingly, downstream phenotypic analyses did not uncover major fitness effects associated with these GRDs, suggesting that a large number of mito-nuclear GRDs may reflect population structure at the mitochondrial level rather than actual genomic incompatibilities. This is further supported by the GRD landscape showing particular large genomic regions associated with a single mitochondrial haplotype. Next, we explored the functional relevance of the detected mitochondrial haplotypes through an association analysis on a set of 259 assembled, non-correlating DGRP phenotypes. We found multiple significant associations with stress- and metabolism-related phenotypes, including food intake in males. We validated the latter observation by reciprocal swapping of mitochondrial genomes from high food intake DGRP lines to low food intake ones. In conclusion, our study uncovered important mitochondrial population structure and haplotype-specific metabolic variation in the DGRP, thus demonstrating the significance of incorporating mitochondrial haplotypes in geno-phenotype relationship studies.

## Introduction

Phenotypic variation is driven by genetic and environmental factors. Genome-wide association (GWA) studies attempt to identify the relationship between genotype and phenotype. However, these studies are often limited by the sole use of nuclear variants^1^ despite the fact that some recent GWA studies have revealed associations between mitochondrial genomic variants and obesity, type 2 diabetes, multiple sclerosis, and schizophrenia in humans^2–5^. The detection of these mitochondrial variants tends to be challenging due to the relatively low extent of mitochondrial genetic variation as compared to the nuclear genome. Other challenges that complicate the detection of genetic determinants (both mitochondrial and nuclear) in human populations involve non-standardized lifestyles, cultural differences and upbringing, and genetic background (population stratification)^6^.

Complementary studies using higher model organism-based genetic reference populations, such as the mouse BXD^7^ or *Drosophila* Genetic Reference (DGRP) panels^8–10^, are in this regard advantageous since phenotyping is performed in a controlled environment and the genetics, or manipulations thereof, are not affected by environmental variation. However, in both these panels, mitochondrial genomic variation has so far been largely ignored. For the BXDs, this is because all lines feature the same maternal-derived mitochondrial haplotype. In contrast, DGRP lines are genetically independent^8,9^, but to our knowledge, no concerted efforts have been devoted toward the extensive characterization of their mitochondrial genomic variation. Indeed, only two studies have so far explored this question, but both suffered from low local sequence coverage issues since they depended on data from whole organism DNA sequencing^11,12^. Moreover, they did not utilize the resulting variant catalogue to explore the impact of mtDNA variants on phenotypic variation. Thus, an in-depth characterization of mitochondrial variation and to which extent it affects phenotypic diversity is still lacking.

In this study, we postulated that a high-resolution catalogue of mitochondrial variants could greatly benefit genotype-to-phenotype relationship studies. This is because a link between mitochondrial and nuclear genomic variation has already been shown to affect metabolic traits in flies^13,14^. For instance, lifespan^14^ and developmental time^15^ were drastically affected by the placement of divergent mitochondria in differing nuclear backgrounds suggesting that the mito-nuclear relationship has the potential to affect a wide-range of other phenotypes as well. While these studies nicely showed the functional importance of mitochondrial and nuclear genomic variant interactions, the mitochondrial genomes used in these studies were highly divergent and the effect of individual (non-deleterious) mitochondrial alleles remains poorly understood. Therefore, characterizing the full extent of mitochondrial (genomic) variation within the DGRP, and integrating this data in GWA analyses could be valuable for the detection of (novel) genetic determinants of phenotypic diversity.

To do so, we assessed mitochondrial variation within the DGRP both at the genotype and phenotype level. First, we produced a high-resolution map of mitochondrial variants for the DGRP by sequencing libraries enriched for the mitochondrial genomic fragments of individual lines. We found substantial genomic variation, at a low allele frequency, in the form of single nucleotide polymorphisms, insertions and deletions, and putative heteroplasmy. These variants allowed for the construction of mitochondrial haplotypes revealing population structure at the mitochondrial level. The link between the mitochondrial and nuclear alleles is distinctly present with the finding of large numbers of genotype ratio distortions (GRDs). These GRDs followed a block-like pattern which rationalize the mitochondrial haplotypes that we identified and thus further support the presence of population structure at the mitochondrial level. Moreover, given that we did not find a biological function, or phenotype, for the complete GRDs that we tested nor in our GRD-phenotype association analyses, we believe that the population structure that we observed is due to an already existing structure upon establishing the DGRP. Finally, we demonstrate that, even though population structure is present, the mitochondrial haplotypes can affect phenotypes. Mitochondrial variation at the haplotype level was significantly associated with 12 metabolic or stress-related phenotypes. As a proof of concept, we selected “food intake in males” and experimentally validated the functional effect of mitochondrial haplotypes. Our study therefore demonstrates the importance of implementing mitochondrial variation in genotype-to-phenotype association studies to reveal novel mitochondrial genetic determinants of quantitative traits, not only in *Drosophila* but also in human GWA studies.

## Methods

### Fly populations and fly rearing

A full list of DGRP lines used in this study can be found in **Supplementary Table S1**. All lines were obtained from the Bloomington Stock Centre (Indiana, USA). Flies were kept at a constant light:dark cycle of 12hr:12hr at 25C and 60-70% humidity. Flies were fed a medium containing: 58.8 g yeast (Springaline BA10), 58.8 g Farigel wheat (Westhove FMZH1), 6.2 g agar powder (ACROS 400400050), 100 ml grape juice (Ramseier), 4.9 ml propionic acid (Sigma P1386), 26.5 ml of methyl 4-hydroxybenzoate (VWR ALFAA14289.0, stock: 400 g/L in 95% ethanol), and 1L water.

### Mitochondrial DNA extraction for mitochondrial DNA enriched libraries

The extraction of mitochondrial DNA consisted of two parts: Mitochondrial extraction and subsequent DNA extraction. First, extraction of mitochondria from 40 fly whole bodies (or 20 mg mouse liver) was based on a protocol from Schwarze et al. 1998^16^. Flies frozen in liquid nitrogen were gently crushed with a plastic pestle in 300 µL mitochondrial isolation medium (MIM; 250 mM sucrose, 10 mM Tris (pH = 7.4), 0.15 mM MgCl_2_). Once the samples were fully homogenized, 700 µL MIM was added while rinsing off any remains on the pestle. Samples were then centrifuged at 850x *g* for 5 minutes at 4°C to reduce debris in the following steps. Supernatant was carefully transferred to a new tube and samples were centrifuged at 1000x *g* for 5 minutes at 4°C. This caused larger cell debris and any chitin remains to precipitate while the remaining supernatant largely consisted of floating mitochondria. Supernatant was then transferred to a new tube and centrifuged at 13,000x *g* for 5 minutes at 4°C followed by washing with 1 mL of MIM. The supernatant was then again transferred to a new tube after which the wash was repeated once more to obtain a pure mitochondrial pellet. If desired, one can store the mitochondrial pellet at −80°C at this point and the supernatant should be discarded. Second, to extract mtDNA from the mitochondrial pellet, 500 µL of RNAse/DNAse-free water was added to resuspend the pellet, followed by adding 500 µL Phenol:Chloroform:Isoamylalcohol (25:24:1; Sigma 77617) and vortexing for 20 seconds. Samples were centrifuged at 14,000 rpm for 5 minutes at room temperature. The upper phase (~500 µL) was then transferred to a new tube and 500 µL Chloroform (Fisher C/4960/PB08) was added. After vortexing for 20 seconds, samples were centrifuged at 14,000 rpm for 5 minutes at room temperature. The upper phase (~500 µL) was transferred to a new tube. Mitochondrial DNA was then precipitated by adding 30 µL of 5M NaCl (0.3M final concentration), 3 µL Glycogen (Thermo Scientific R0561), 530 µL Isopropanol (=1V) and gently inverting the sample six times. Next, precipitation of mtDNA in the samples either took place on ice for 30 minutes, or overnight at −20°C with the latter providing the greatest yield. Following incubation, samples were centrifuged at 14,000 rpm at 4°C for 60 minutes. The mitochondrial DNA pellet was washed once with 1 mL 70% ethanol, eluted in 30 µL RNAse/DNAse-free water, and stored at −80°C until use.

### Library preparation and sequencing of mitochondrial DNA-enriched libraries

Prior to library preparation, mitochondrial DNA samples were quantified by either the Qubit dsDNA HS assay kit (Invitrogen Q32851) or the Quant-iT PicoGreen dsDNA assay kit (Invitrogen P11496). Samples were accordingly diluted to 5 ng/µL and quantified again to ensure the correct concentration post-dilution. For each library, 5 ng of input mtDNA was diluted in a total volume of 11.77 µL. To each well, 8.23 µL of a master mix was added consisting of 4 µL 5X TAPS-MG (50 mM TAPS (Sigma T5130), 25 mM MgCl_2_ (Sigma M2670)), 4 µL 40% PEG 8K (adjusted to RT prior to incubation; Sigma P5413), and 0.23 µL of 11 µM in-house generated Tn5 Transposase^17^. Tagmentation of the samples was done by incubation at 55°C for 4 minutes and samples were immediately placed on ice for 2 minutes afterwards. The reaction can be stopped by either adding RNAse/DNAse-free water or by using the Zymo Research gDNA Clean-Up & Concentration kit (Zymo Research D4066) following manufacturer’s instructions using the DNA Binding Buffer ratio of 5:1. We found that using a DNA clean-up kit resulted in the most consistent results and is likely due to the potential residues of organic DNA extractions that may interfere in PCR reactions. Samples were eluted with 25 µL Elution Buffer. Amplification of the libraries was done by adding 25 µL 2X NEB Next High-Fidelity Buffer (NEB M0541S), 2.5 µL of i7 and 2.5 µL of i5 barcoded primers. Conditions for the PCR were as follows: 1) one cycle at 72°C for 3 minutes followed by a single cycle at 95°C for 30 seconds; 2) 12-15 cycles of 10 seconds at 95°C, 30 seconds at 55°C, 1 minute at 72°C; 3) One cycle for 5 minutes at 72°C. Libraries were purified by using AMPure XP magnet bead selection (Beckman Coulter A63881) at a 1X ratio. Briefly, 55 µL of AMPure XP beads were added to 55 µL of library sample and gently mixed by pipetting 10 times. Samples were incubated for 10 minutes at room temperature before placing them on a magnetic holder and incubating another 10 minutes at room temperature. Supernatant was removed, and libraries were washed twice with 200 µL freshly prepared 80% ethanol. After washing, samples were left to air dry for 10 minutes and subsequently removed from the magnetic holder. The mitochondrial DNA enriched libraries were then eluted by adding 23.5 µL elution buffer and carefully pipetting the mixture 10 times and allowing this to incubate for 5 minutes. Finally, samples were placed again on the magnetic holder and incubated for 5 minutes after which 22 µL of mitochondrial DNA enriched libraries were transferred to new tubes. Libraries were quantified by either Qubit dsDNA HS Assay Kit (Invitrogen Q32851) or Quant-iT PicoGreen dsDNA assay kit (Invitrogen P11496). Size distribution of the libraries was assessed by using the High Sensitivity NGS Fragment Analysis Kit (Advanced Analytical DNF-474). Libraries were sequenced in three different batches of which two used an Illumina HiSeq-2500 (paired-end; 100 cycles, loaded molarity 5 nM) and one used an Illumina NextSeq 500 (single-end; 75 cycles, loaded molarity 1.8 pM). Accession numbers for the datasets can be found in **Supplementary Table S2.**

### Assessing enrichment of mitochondrial DNA over nuclear genomic DNA

We assessed the enrichment of raw mtDNA or mtDNA fragments in our libraries via qPCR using primers for four nuclear loci and four mitochondrial loci. Each qPCR reaction contained 200 µM of the primer mix (Forward and Reverse, see **Supplementary Table S3** for a list of primers), 1.5 µL of diluted sample (1:20), and 5 µL of 2X SybrGreen PowerUp Mastermix (Invitrogen A25743). Samples were measured either on a QuantStudio 6 (Applied Biosystems) or StepOnePlus Real-Time PCR system (Applied Biosystems) for 384-well and 96-well formats, respectively. Enrichment was then calculated using the 2^−∆∆CT^ method. All measured samples were compared to a single reference sample of 1 ng/µL genomic DNA from the *iso-1* genotype and one of the four nuclear loci.

### Data pre-processing and genotyping

While our method enriches for mitochondrial DNA, the presence of nuclear DNA cannot be entirely prevented. As such, we hypothesized that various nuclear regions could still be used to ‘genotype’ the sequenced samples and correct for potential swapping. A schematic overview of the sequence data pre-processing, genotyping, and variant calling pipeline is provided in **Supplementary Figure S1**. Raw reads obtained from the Illumina HiSeq-2500 and the Illumina NextSeq 500 were first demultiplexed with BRB-seq tools v1.1 (https://github.com/DeplanckeLab/BRB-seqTools^18^) and trimmed with Trim Galore v0.4.4^19^. For each sample, the paired-end or single-end mode of sequencing was taken into account when trimming and in further downstream steps. Trimmed reads were first mapped to dm3(UCSC, BDGP R5, Apr 2006) with BWA mem v0.7.13^20^ followed by duplicate read removal with Picard v2.2.1 (http://broadinstitute.github.io/picard) using default settings. Mapping statistics (i.e. number of mapped and duplicated reads) for each sample were accessed via SamTools v1.3^21^. We then used GATK v3.6-0^22^ to perform local realignment around indels followed by HaplotypeCaller in GVCF mode (–emitRefConfidence GVCF) to only call nuclear variants with the minimum phred-scaled confidence threshold set at 30 and the emission confidence threshold set at 10. For every sample, GATK SelectVariants was used to filter out all indels and MNPs, and SNPs with a depth of coverage less than 5X from the overall set of variants. The remaining set of variants (SNPs) was then used as input for GATK GenotypeGVCFs for the genotyping of all samples as a cohort applying the same phred-scaled confidence threshold and emission confidence threshold used in HaplotypeCaller. Finally, only bi-allelic SNPs with a Fisher strand score of FS > 30.0 and quality by depth QD < 2.0 were selected for the comparison with the DGRP2 reference VCF^8,9^. For each DGRP sample, we assessed the top three of genotype matches. If the tested DGRP and the expected DGRP were the highest ranked, had a >90% match and the second and third match were at least 5% lower, we considered it as a clean match. In case the first ranked expected DGRP did not match the tested DGRP, however, and >90% of the tested loci matched the expected DGRP, and the second and third matches were 5% lower, then we considered this a mislabelling and renamed the DGRP accordingly. Notably, in most of the samples (94%), more than 99% detected SNPs were found matching to the corresponding DGRP line.

### Mapping and mitochondrial variants calling

The overall pipeline (see **Supplementary Figure S1**) to call mitochondrial variants is similar to calling nuclear variants, with the main exception being that trimmed reads were mapped to dm6(UCSC, BDGP R6 + ISO-1 MT (NC_024511.2), Aug 2014). This genome version provided a significant improvement in alignment quality for the mitochondrial genome over version dm3. Parameters for each package remained unchanged unless noted differently. Variant calling was performed using GATK HaplotypeCaller including both nuclear and mitochondrial variants. GATK SelectVariants was then used to restrict to mitochondrial variants only after which the GVCFs were merged and another run of variant calling on the mitochondrial genome was performed using GenotypeGVCFs. High quality variants were selected by filtering out variants with depth of coverage less than 10 (DP < 10), Fisher score more than 60 (FS > 60.0) and RMS Mapping Quality less than 10 (MQ < 10). Finally, we only retained variants within the main coding part of the mitochondrial genome (up to 14,917 bp) due to the inaccuracy in mapping short reads to the 4.5 kb AT-rich repeat region. The resulting set of mitochondrial variants was annotated with snpEff v4.2^23^ using the following parameters: -no-downstream -no-upstream -no-utr. Overall statistics on the number of reads and coverage per sample can be found in **Supplementary Table S4**. Randomly selected variants were verified via Sanger sequencing of multiple loci for particular DGRP lines (**Supplementary Table S5-S6**).

### Detection of nuclear encoded mitochondrial DNA fragments (NUMTs)

We assessed the presence of mitochondrial fragments integrated in the nuclear genome (NUMTs) using methods described earlier^24^. First, using our mouse mitochondrial sequence data, the putative location of NUMTs was detected by aligning the mouse mitochondrial reference genome (mm10, UCSC, GRCm38, Dec 2011) to mouse nuclear genome with NCBI-blast v2.2.28. All hits with a length >50 bp were kept as potential NUMT sites and considered as a NUMT genome. Reads were then aligned to the NUMT genome using BWA v0.7.13 and duplicates were removed with picard v2.2.1. Reads with a perfect match to the NUMT genome were aligned to the mitochondrial genome. Reads that did not show a perfect match to the mitochondrial genome were considered to be contaminations from NUMTs. We applied the same method to our fly mitochondrial sequence data.

### Comparison with the set of mitochondrial variants from Richardson et al. 2012

Previously, Richardson et al. (2012)^11^ analysed the DGRP sequence data from Mackay et al. (2012)^8^ for mitochondrial variants. We compared our findings by running our data through the variant calling pipeline described in Richardson et al. (2012)^11^ using the pipeline and reference genome available at that time (dm3: chrU, ch3L 10Mb-11.2 Mb and *Wolbachia*) as our variant calling pipeline is less compatible with the type of sequence data obtained from Mackay et al. (2012)^8^. We restricted the two datasets to the 134 overlapping lines between our study and Richardson et al. (2012)^11^. We used the variant coordinates identified by Richardson et al. (2012)^11^ in “Dataset S4” for our final comparison.

### Resolving the intergenic repeat region between mt:ND3 and mt:tRNA:A

The region from 5,959 bp to 5,983 bp between mitochondrial genes mt:ND3 and mt:tRNA:A contains a highly heteroplasmic AT-repeat region. We confirmed the existence of this repeat region via Sanger sequencing of various DGRP lines for this region. In order to fully resolve this region for all DGRP populations, we first selected reads overlapping at least one bp between 5,959-5,983 bp for each sample using bedtools intersect v2.25.0^25^. Our previous mapping of the surrounding regions and results from the Sanger Sequencing already provided us with a conserved pattern flanking this region (5’-CTA[repeat]GGG-‘3). Thus, from our pre-selected reads, reads were extracted that spanned both the 5’- and -‘3 region. If >10 unique reads provided a single pattern, we considered this pattern to be true.

### Detection of putatively heteroplasmic loci

To detect putative heteroplasmy from high-throughput sequence data, we used a method described before^26^. Briefly, reads with Phred quality score <23 and with any 5 neighbouring base pairs (both directions) with Phred quality <15 were discarded. To consider loci heteroplasmic, we followed a set of criteria that was applied per nucleotide position. First, the depth of coverage was at least 20. Second, the minor allele was present in at least 15% of the reads. And third, the minor allele should be present in at least two reads of each strand. The heteroplasmic detection does not rely on previously annotated vcf files. Therefore, it may detect variants or putative heteroplasmic loci that were not picked up by our variant calling pipeline. These non-overlapping loci were removed from our analysis leaving only those loci for which we had previously detected a variant.

### Verifying mitochondrial variants via Sanger sequencing

To confirm our results from the variant calling pipeline, we used Sanger sequencing of various selected loci. Per locus, we selected reference strains (i.e. *w^1118^*) and DGRP lines that had a mix of reference and alternate alleles between each other. For each sample, we used 100 ng of regular genomic DNA from five females which was amplified in a volume of 20 µL consisting of 10 µL 2X NEB Next High-Fidelity Buffer (NEB M0541S), 1 µL of 10 µM primer mix (forward and reverse, see **Supplementary Table S5**) and RNAse-DNAse free water. All samples were sequenced with the M13 forward primer. Sequence results were aligned and analysed using MEGA v7.021^27^.

### Population structure analyses

For the identification of haplotypes, we performed multiple sequence alignment between all the strains using MAFFT^28^. Finally, the haplotype network was inferred using the TCS software^12,29^. To further define the remaining surrounding clusters, we used a *k*-means clustering approach to accurately define haplotypes. However, regardless of the chosen alpha, no clear *k* could accurately be selected. Therefore, using the TCS-based network as a framework, we continued with manual annotation of haplotypes by considering a group of lines to form a haplotype when a particular variant or set of variants are shared by >5 DGRP lines but not by the central haplotype or another haplotype. Lines that would cluster with other non-DGRP reference strains yet did not meet the criteria of >5 DGRP lines, would still be considered to be a haplotype.

To determine the degree of similarity or differentiation between the observed haplotypes, we assessed the genetic distance between clusters using a *G_ST_* estimator (Hedrick’s *G’_ST_*). We used the multiple fasta aligned sequences (used for the MAFFT alignment) which were imported in ‘R’ using apex (v1.03). Subsequently, we set the population strata based on our pre-defined haplotypes and calculated the pairwise *G’_ST_* for each pair of haplotypes. To test for significance, we bootstrapped the dataset and recalculated the pairwise *G’_ST_*. Finally, we performed permutations on the data by reshuffling the haplotype associated DGRP lines and recalculating the *G’_ST_* estimators.

Furthermore, we only considered the first variant of a set of linked variants (clusters) for downstream association analyses to reduce the number of mitochondrial variants and thus statistical tests. This resulted in a set of 12 haplotype defining variants and three variants that were more widespread among the DGRP lines (see also **Figure 3d, Supplementary Table S7-S8**).

### Genotype Ratio Distortions (GRDs)

To infer genotype ratio distortions (GRDs) between the mitochondrial and nuclear genomes, we applied the method described in ^30^. In our primary GRD analysis, we restricted the set of *mitochondrial* variants to bi-allelic non-MNP variants with a minor allele frequency >0.05 and for which at least 150 DGRP lines have a confident variant call using vcftools v0.1.14 (–min-alleles 2 –max-alleles 2 –maf 0.05 –max-missing 0.88)^31^. Moreover, to prevent the inflation of the number of statistical tests, we reduced our set of mitochondrial variants to haplotype-defining variants and three remaining variants representing unique clusters (see **Population structure** section in **Methods**). Heterozygous variants were not used and marked as missing data. After applying these filters, a set of 12 mitochondrial variants remained which were used for further downstream analysis (**Supplementary Table S8**). The set of *nuclear* variants were selected by converting the dm3-based DGRP2 VCF^8,9^ to dm6 via CrossMap v0.2.5^32^. Applying the same filters as for the mitochondrial variants resulted in 1,722,758 nuclear variants that were used for the analysis. We further restricted the calculations to include only those pairs of variants for which at least 150 DGRP lines had confident variant calls for a given mitochondrial and nuclear variant combination. For each combination of mitochondrial and nuclear variants, a χ^2^-test was performed. We adjusted p-values to correct for multiple testing according to a 4-step procedure described in section 7 by Benjamini and Bogomolov (2014)^33^. First, we calculated a p-value of a χ^2^-test for each of 1,722,758 x 12 mito-nuclear variant combinations. Second, the intersection hypothesis was computed as Simes test for every tested nuclear variant using set of 12 *p*-values. Second, we applied a Benjamini-Hochberg (FDR) procedure to all the 1,722,758 p-values calculated at previous step and selected 14,643 p-values with an FDR < 0.1. Then, we selected 14,643 nuclear variants for which the *p*-values passed the threshold and applied a Benjamini-Hochberg correction (FDR) procedure to all raw, computed *p*-values at step 1 (χ^2^-test *p*-values (12 x 14,463)). Fourth, a selection was made between mito-nuclear variant pairs for which the FDR computed in the previous step was less than the nominal FDR cut-off (0.1 / 1,722,758) multiplied by the number of the selected mito-nuclear variant pairs from the previous step (14,463). The resulting cut-off was then 8.5 x 10^4^. Finally, in line with Corbett-Detig et al. 2013^30^, we only considered mito-nuclear GRDs that contained at least one putative significant GRD on either flanking side within a 50 bp range (see **Supplementary Table S9-S10**).

### Selection and crossing of populations to assess mito-nuclear incompatibility

To empirically assess whether complete genotype ratio distortions reflect mito-nuclear incompatibility, we selected DGRP lines following a set of parameters to test experimentally. First, we selected lines that contained the alternative alleles for the linked nuclear variants 2L_8955411_SNP, 2L_8955426_SNP, and 2L_8955438_SNP which are located in the gene *CG31886* and the reference allele for X_9007384_SNP located in *rdgA*. These lines were further restricted to the ones with the alternative alleles for mitochondrial variants chrM:2349_SNP, chrM:5500_SNP, chrM:11128_SNP, chrM:12682_INDEL, and chrM:12898_SNP. We dubbed this configuration ‘AA’, the first letter indicating the nuclear allele (alternate, from the *CG31886* perspective) and the second the mitochondrial allele (also alternate). Second, to reduce effects of genomic inversions, we selected those lines without inversions. Third, we were further restricted to select populations containing *Wolbachia* given the limitations by the previous criteria. Next, we selected lines with the ‘RR’ configuration (= reference nuclear (CG31886 perspective) and reference mitochondrial alleles). Again, only lines were selected that were infected with *Wolbachia* and had no genomic inversions. To mitigate the effects of other mitochondrial variants, we used populations with similar mitochondrial haplotypes. Therefore, we selected lines from the consensus mitochondrial haplotype directly or those only differing one mitochondrial variant from the consensus mitochondrial haplotype (‘Central MH’, see methods below). Finally, applying these selection criteria, we selected the following lines with the AA configuration: *DGRP_189*, *DGRP_801*, *DGRP_882*. And for the RR configuration: *DGRP_153*, *DGRP_287*, *DGRP_306* where *DGRP_287* was selected from lines that differ only one mitochondrial variant from the consensus mitochondrial haplotype (see **Supplemental Table S11** and **Supplemental Figure S2** for the crossing scheme).

### Development assay for the mito-nuclear complete GRD populations

To test if mito-nuclear incompatibility was responsible for the complete *CG31886*/*rdgA*-GRD, we assessed the impact on *Drosophila* development. We measured the development time from egg to adult on hybrid-F2 and on flies that were further backcrossed to F10 populations (see **Supplemental Figure S2** for the crossing scheme). The F2 generation is the first generation in which 50% of the offspring have the homozygous reference nuclear variant (*CG31886* perspective) combined with the alternative mitochondrial variants (the ‘RA’ configuration). We cannot exclude the potential influence of genetic interactions that may occur from other loci and we assume that these would be wide-spread throughout the genome and throughout the offspring. For the collection of eggs of each population, we used demography cages containing 75 female F1 hybrid offspring from AA females and RR males and crossed these to 75 RR males. As a control, we backcrossed 75 male F1 hybrid offspring from AA females and RR males with 75 RR females. On the bottom of each demography cage, 60 mm petri dishes containing grapefruit juice agar plates were mounted on which a small amount of fresh yeast paste was placed on the agar. Plates and yeast paste were replaced daily, and on the third day, eggs were collected in two-hour intervals. Development time was adjusted accordingly. Per cross, three replicates were used containing 50 eggs per replicate. We used a blocking scheme to prevent potential confounding effects from the placement of the replicates. We applied a similar setup for the measurement of the development time of the F10 generations although we did not cross these to AA lines after the initial F10 emerged. Statistics for the development time were analysed using SAS JMP Statistics v9.

### Climbing activity for mito-nuclear complete GRD populations

To assess the climbing ability of the *CG31886*/*rdgA*-GRD lines, we used 4- to 5-day old females of the F10 generation. Per population, we used three biological replicates, each containing 10 females. Sorting of the flies under CO2 anaesthesia was done 22-24 hours prior to the experiment to avoid the effects of the anaesthetics on the experiment. On the day of the experiment, tubes with flies were mounted on a *Drosoflipper* rack (http://drosoflipper.com), which allows for the measurement of 10 tubes at the same time. Prior to the experiment, all flies were transferred to empty tubes marked at a specific height (7 cm) and were mounted on the other side of the *Drosoflipper* rack. The *Drosoflipper* rack with the flies was then tapped, after which the climbing of the flies was recorded. Scoring was done by calculating the fraction of flies in the tube that were able to pass the marked line 10 seconds after the tapping stopped. Technical replicates were repeated at 1.5-minute intervals. Positioning of the tubes was randomized so that no genotype was always placed at the same location in the *Drosoflipper* rack. Experiments were performed between 08:30 and 10:00 AM (Light cycles from 08:00-20:00).

### GRD-phenotype association analysis

To uncover the potential relationship between genotype ratio distortions (GRDs) and fitness phenotypes, we first assembled a compendium of >600 publicly accessible DGRP phenotypes (Bevers *et al*., *in prep*.) and selected 76 phenotypes that we considered as putative proxy read-outs of “fitness” after which we further reduced this list to 29 independent, uncorrelated fitness phenotypes (see **Supplementary Table S12** for the source). The following steps will therefore be applied to the set of 29 uncorrelated fitness phenotypes. For each of the 1,845 detected GRDs, we assessed the enrichment of a given mito-nuclear allelic combination in the top and bottom 20% of a given fitness phenotype using a hypergeometric test, resulting in eight tests per GRD per fitness phenotype. We restricted our analysis to only those GRD-phenotype pairs in which 100 DGRP lines were genotyped for both the mitochondrial and nuclear variants, and which had a phenotypic measurement.

To perform multiple testing correction, we applied the 4-step procedure developed by Benjamini and Bogomolov (2014) to each of the eight sets of tested mito-nuclear genetic combinations and phenotypes. First, for every GRD, we calculated the intersection hypothesis via a Simes test on 29 p-values (from each phenotype) obtained from the hypergeometric test. Second, we applied a Benjamini-Hochberg (FDR) procedure to all the 1,845 *p*-values calculated at the previous step and selected adjusted *p*-values with an FDR < 0.2. Out of the eight sets of tested mito-nuclear allelic combinations and phenotypes, only the (mito)ALT-(nucl)REF combination yielded three significant *p*-values for chrM:7424-chr3R_24564153, chrM:7424-chrX_14792890, chrM:7424-chrX_14792895 passing that threshold (raw *p*-values: 1.2e-06, 1.72e-06, 1.72e-06 respectively). Third, we selected raw p-values calculated at step 1 for the three mito-nuclear allelic combinations (in (mito)ALT-(nucl)REF configuration only) mentioned at the previous step, yielding a matrix of 3 x 29 p-values to which we applied a FDR < 0.2 correction to. The p-value cut-off for the final stage was obtained by multiplying the nominal FDR cut-off (0.2 / 1845) with the number of selected mito-nuclear allelic combinations from step 3. This is done in order to control for the FDR on both GRD and phenotype levels in accordance with the first theorem in Benjamini and Bogomolov (2014).

### Mitochondrial haplotype association analysis

For a mitochondrial genome-wide association analysis, we would be unable to use most of the mitochondrial variants that we detected because of the low allele frequency and pruning of the remainder of the variants resulted in the haplotype-defining variants. We therefore used the haplotypes for our association analysis. We again employed the assembled compendium of >600 publicly accessible DGRP phenotypes (Bevers et al., *in prep*.) which was reduced to 254 uncorrelated phenotype groups detected via hierarchical clustering. Subsequently, the association analysis was performed in two parts. First, to reduce the impact of multiple test corrections, we performed an ANOVA on each phenotype to screen for signatures of variation using a relaxed *p*-value cut-off of 0.3. This resulted in 12 phenotypes in which we may detect variation. For each of these 12 phenotypes, we employed a Tukey’s HSD test to detect significant differences (*p* < 0.05) between the haplotype pairs (see **Supplementary Table S13**).

### Assessment of feeding behaviour of *Drosophila*

Measuring the feeding behaviour of *Drosophila* was done using the CAFE method^34^ with a few adjustments, such as the fact that we exposed the flies to a 2-hour starvation period (instead of an 18-hour period). In our hands, various extreme food intake lines did not survive the starvation period or were more susceptible to it (data not shown). After this adjustment, results of Garlapow et al. (2015) ^34^ and ours correlated in their pattern. Furthermore, we used five biological replicates per genotype or cross, each consisting of six 5- to 7-day old male flies that were placed in a tube containing 5 mL of 1.5% agarose medium. Two 5 µL capillaries (Aldrich BR708707) filled with a freshly prepared solution composed of 4% sucrose, and 0.1% (w/v) erioglaucine disodium salt (Sigma 861146) were placed in each vial 2 hours after starvation (at 10:00 AM). The starting position of the solution was marked prior to the placement of the capillaries in the vials. We also placed capillaries in a tube with the agar solution but without flies which would be used to correct for evaporation of our sucrose solution^34^. Flies were allowed to feed for 10 hours at which point the capillaries were removed and imaged for further scoring. Finally, we analysed the food intake using ImageJ by measuring the total length up to the mark and subtracting the length of the leftover blue sucrose solution. Food intake was corrected by the number of flies left at the end of the assay and by the evaporation that occurred in the empty vials.

## Results

### A robust mitochondrial DNA enrichment workflow

To achieve high mitochondrial DNA (mtDNA) sequencing coverage with the aim of increasing the overall confidence in variant calling, we developed a new approach for preparing mtDNA-enriched sequencing libraries. Briefly, building on an existing mitochondrial extraction method^16^, we used differential centrifugation to isolate mitochondria from 40x whole body *w^1118^* flies (**Figure 1a**). For the extraction of mtDNA from the previous step, we used a phenol:chloroform:isoamylalcohol extraction method and isopropanol precipitation^35^. To assess the efficacy of our method, we measured the amount of mtDNA fragments versus nuDNA fragments in mitochondria-enriched samples and whole organism DNA samples via qPCR, revealing high mtDNA enrichment (**Figure 1b**). Subsequently, we tagmented the mtDNA with an in-house produced Tn5 transposase^17^ to produce mtDNA-enriched sequencing libraries. After amplification of the libraries, the enrichment of mtDNA was still present and nuclear genomic regions were not preferentially tagmented or over-amplified during the library preparation step (**Figure 1b**).

**Figure 1.**
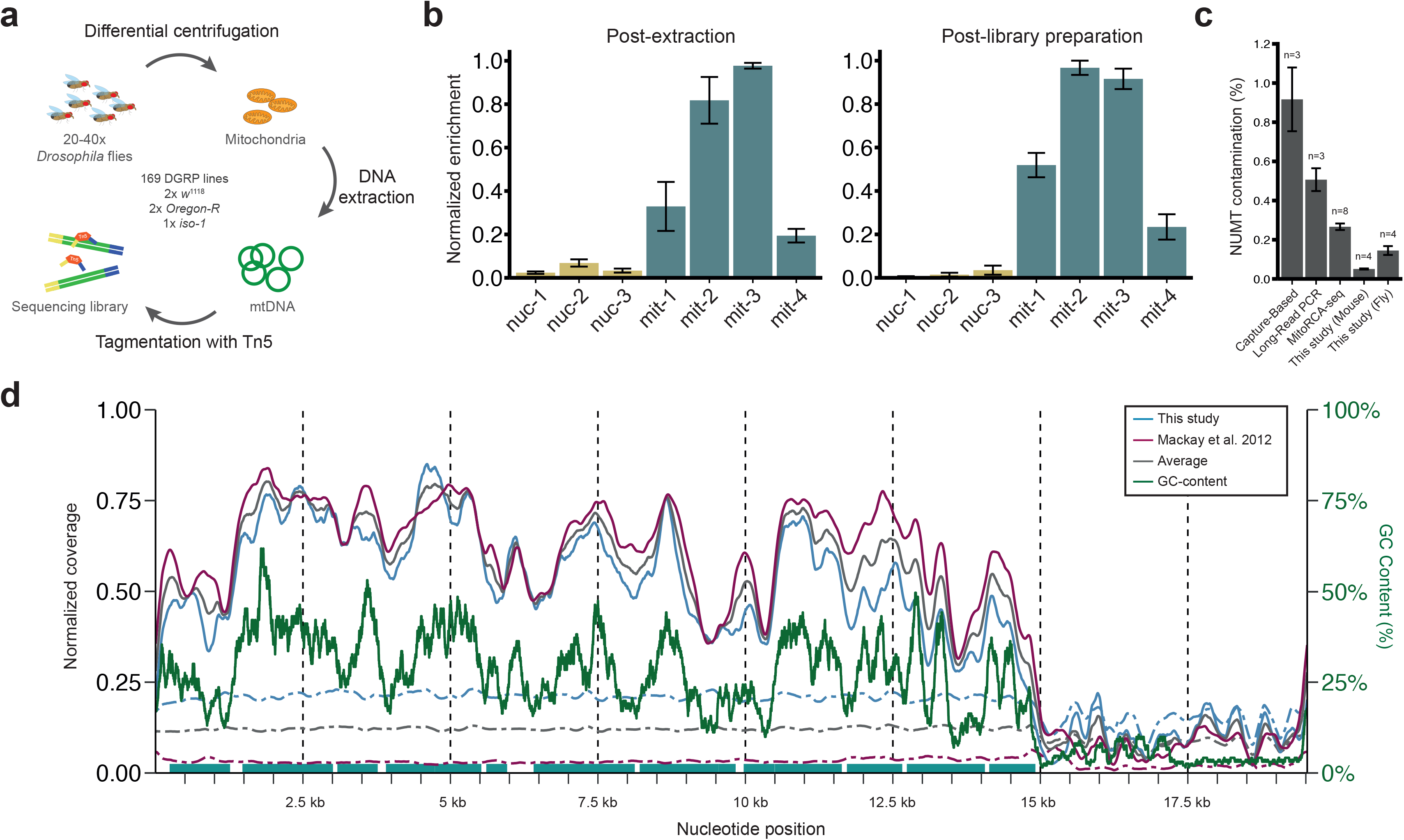
Sequencing of mitochondrial DNA-enriched libraries for 169 *Drosophila* Genetic Reference Panel (DGRP) lines. **a)** Overview of the newly developed protocol for preparing mtDNA-enriched sequencing libraries using tagmentation. **b)** Normalized enrichment of mtDNA in samples post-DNA extraction (left) and post-library preparation (right). For the analysis, four different mitochondrial (green) and nuclear (yellow) loci were amplified. One of the nuclear loci was used as a reference. **c)** Comparison of the percentage of contamination by nuclear mitochondrial fragments (NUMTs) for different mtDNA sequencing methods. Data for Capture-Based, Long-Read PCR and MitoRCA-seq were used from Ni et al. 2015^24^. **d)** Comparison of the normalized coverage between our mtDNA sequencing method (blue) and the regular sequencing profile of Mackay et al. 2012^8^ (pink) across the mitochondrial genome. Solid lines depict the coverage profile per DGRP line and dashed lines depict the overall coverage per bp achieved by each of the studies. GC-content is depicted with a green solid line (200 bp bins). Light green blocks represent the mitochondrial genes excluding tRNAs.

A particular concern for the detection of mitochondrial variants is the potential contamination of nuclear genome-encoded mitochondrial genes or fragments (NUMTs). To assess the purity of obtaining mtDNA-only fragments, we compared the presence of NUMTs in our sequence data to the presence of NUMTs in public datasets. To our knowledge, there is no mtDNA-enriched sequence data available for *Drosophila melanogaster*. Therefore, in parallel, we extracted mtDNA from C57BL/6J mouse liver and prepared libraries in a similar fashion to infer the potential and effect of our method on the presence of NUMTs. We compared our method to three existing mtDNA sequencing methods for mouse liver samples^24^ and observed a significantly lower percentage of reads mapping to NUMTs using our approach. This shows the efficacy of our approach in yielding highly pure mtDNA (**Figure 1c**).

The robustness of our mtDNA enrichment approach on both *Drosophila* and mouse samples persuaded us to employ this technique for the whole set of DGRP lines that was available, consisting of 169 lines. In addition, we generated replicate samples for the common reference lines *iso-1*, *Oregon-R*, and *w^1118^*. We obtained a median coverage per sample of 772X (min: 31X, max 4071X, **Supplementary Figure S3**).

Since no *Drosophila* mtDNA sequencing dataset is to our knowledge currently available, we compared our sequencing coverage to the initial DGRP sequence data^8^. This dataset was derived from whole fly DNA sequencing of all DGRP lines with 134 of these overlapping with our study. Considering the mtDNA coverage profile per sample (**Figure 1d**), we found that the normalized mitochondrial genomic coverage of the two studies is comparable. In both our datasets, similar drops in coverage can be observed, and most strikingly in the 5 kb AT-rich repeat region, which is likely driven by GC-content fluctuations in the mitochondrial genome, given the strong (Spearman r^2^ = 0.802 [This study] and 0.889 [Mackay et al. 2012]^8^) correlation between this parameter and normalized coverage. However, and importantly, our dataset yielded on average a more than six-fold greater coverage per bp across the accessible mitochondrial genome (**Figure 1d**). This is further accentuated within the raw coverage of overlapping lines where in our data, 13 lines have loci with a coverage <10X with a median of 61 loci per line being affected versus 43 lines with a median of 256 loci in the Mackay et al. 2012^8^ data (**Supplementary Table S14**). Considering that we multiplexed on average 70 samples per sequence run, this further illustrates the efficacy of our newly described mtDNA isolation approach in robustly enriching for sequencing-compatible mtDNA while minimizing contamination of nuclear genome-derived sequences, including NUMTs.

### A comprehensive DGRP mtDNA variant catalogue

Next, using the GATK Gold Standard pipeline (2016), we identified mitochondrial variants for 169 DGRP lines and the 3 commonly used reference lines (**Figure 2a**, and **Supplementary Figure S1** for the pipeline overview). We focused on detecting variants in the coding region (bp 1 to 14,917) of the mitochondrial genome because of limited coverage of the AT-rich repeat region. Overall, we detected 231 DGRP specific variants yielding ~1 variant every 65 bp (**Figure 2a, Supplementary Table S7+S15**), with individual populations having on average 22 variants (~1 variant per 680 bp per DGRP line, **Supplementary Figure S4**). We found that the majority (91%) of mitochondrial genomic variants within the DGRP consists of single nucleotide polymorphisms (SNPs) with each line also containing at least one multiple nucleotide polymorphism (MNP) (see **Figure 2b**). Furthermore, out of 169 DGRP lines, 161 contained an insertion or deletion (indel). While the majority of indels and MNPs occurred in two or more DGRP lines, most of the detected SNPs had a low minor allele frequency (MAF < 1%) and were in fact unique for a given line (**Figure 2c**). It is unlikely that these unique variants were affected by low coverage or sequencing errors as we found a median coverage of 892X (min 15X, max 3871X, mean 1132X) for these line-specific variants. Moreover, of the 22 variants that we verified using Sanger sequencing, we reached a 100% accordance while two variants were not detected in our mitochondrial DNA enriched sequencing data (**Supplementary Table S6**). Since the accessible mitochondrial genome largely consists of coding regions, we evaluated whether the variants have a potential functional impact or whether these mostly lead to synonymous substitutions (**Figure 2a** inner panel). Nearly half (49%) of the SNPs that were not located within a tRNA or ribosomal RNA were missense variants. In general, synonymous variants had a higher minor allele frequency than missense variants (0.027 vs. 0.009, respectively) suggesting that DGRP line-specific SNPs are likely missense variants. At the gene level, especially *mt:ATPase6* (80%) and *mt:Cyt-b* (67%) were proportionally more affected by the presence of missense variants (amino-acid replacements, see **Figure 2d**).

**Figure 2.**
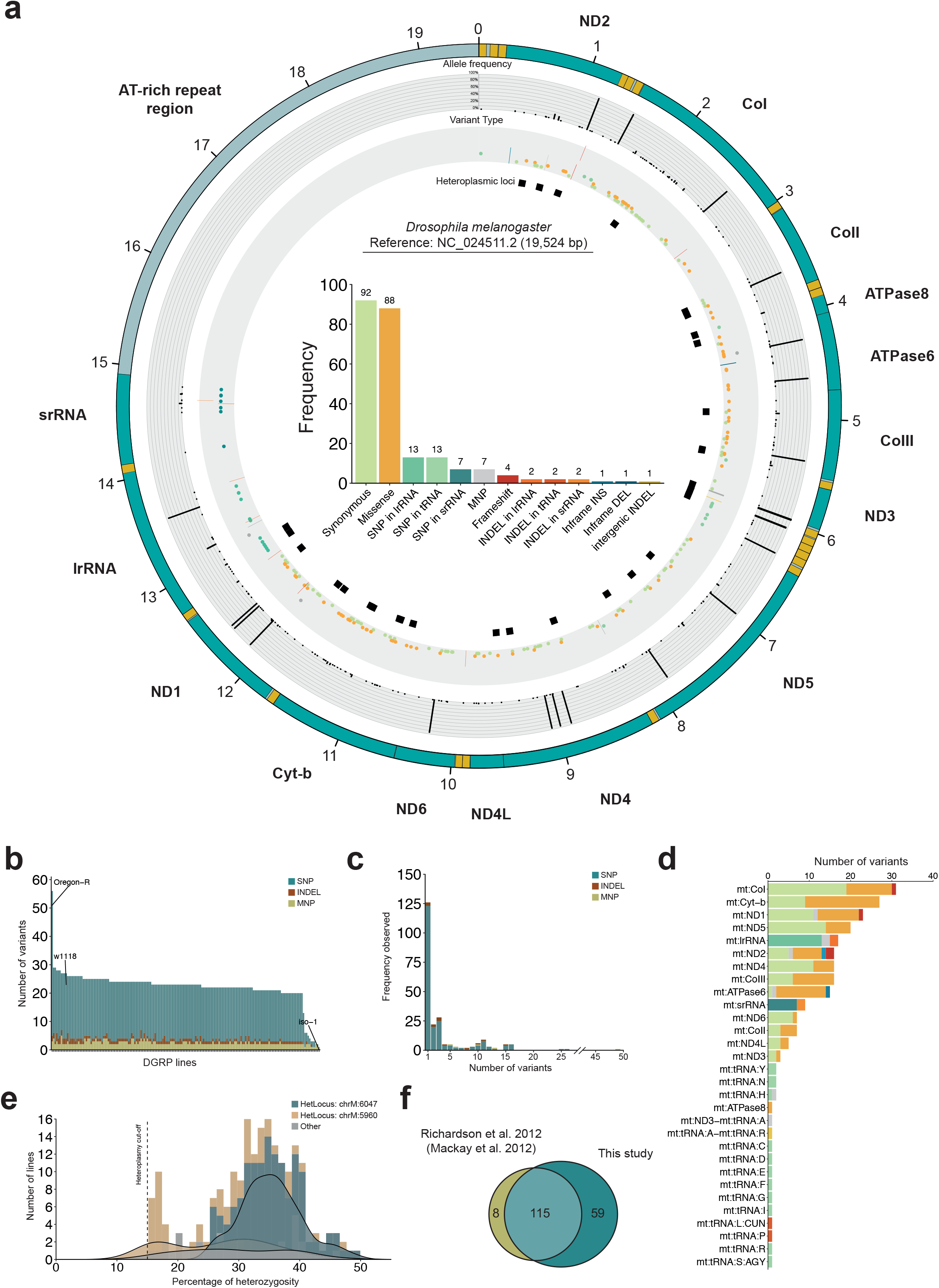
Mitochondrial genomic variation within the *Drosophila* Genetic Reference Panel (DGRP). **a)** Circos plot depicting overall mitochondrial genomic variation. From outer to inner rings: i) gene position, ii) variant allele frequency, iii) variant type (lines are indels), and iv) putatively heteroplasmic variants. The inner bar plot shows the frequency of each variant type. Colours correspond to the ones used in the variant type ring. **b)** Bar plot displaying the number of variants per population. SNPs are in green, indels in brown, MNPs in yellow. **c)** Variant frequency. SNPs are in green, indels in brown, MNPs in yellow. **d)** Variant distribution per gene. Colours correspond to the ones used in the bar plot in **2a**. **e)** Percentage of heterozygosity (a proxy for heteroplasmy) in populations (x-axis). On the y-axis, the number of lines is presented that have a particular putatively heteroplasmic locus. The heteroplasmy cut-off was set at 15%. In green, the heteroplasmic region at chrM:6047, in brown the heteroplasmic region at chrM:5960, and in grey are the remaining heteroplasmic regions pooled together. **f)** Venn diagram of mitochondrial variants detected by Richardson et al. 2012 ^11^ and our study.

Heteroplasmic sites in the mitochondrial genome of *Drosophila melanogaster* have been observed after ~200 generations of mutation accumulation^36^. Within the DGRP, we identified 32 heterozygous variants in 140 lines, reflecting either within population segregation or true heteroplasmic sites, and 92 lines had only one such heterozygous variant. The majority of these 32 variants (29) were only observed once spread over 19 lines, another was present in two DGRP lines, while there were two common variants, both being intergenic. The first, chrM:6047, is an intergenic deletion of 5 bp located between *mt:tRNA:A* and *mt:tRNA:R*. We found that this intergenic deletion is present in 120 lines with a median of 35% of the reads for that site being the alternative variant (**Figure 2e**). The second, putatively heteroplasmic variant resided at chrM:5960 between *mt:ND3* and *mt:tRNA:A*. This locus was picked up during our variant calling because the multiple (>2) alternative AT-repeat alleles were particularly challenging for read mapping due to their repetitive nature. We therefore specifically targeted individual reads containing both the flanking sequences of this locus to accurately resolve the variants within this region (**Supplementary Figure S5a**). We identified that this putatively heteroplasmic intergenic indel followed a distinct pattern of [AT]_x3_[T]_x1_[TA]_x8_ in the reference genome with the largest variant among the DGRP lines showing a TA-tail containing 12 repeats. Moreover, this intergenic region contained substantially more alternative types detecting 15 unique alternative variant types, with each line having a mean of 1.4 types supported by >10 reads. We verified this locus via Sanger sequencing for randomly selected DGRP lines and two reference lines and found that the identified type via Sanger seq, is also the type with the maximum number of reads for that particular line (**Supplementary Figure S5b, Supplementary Table S6**). When we explored the distribution of the types of intergenic regions, most of the DGRP lines did not contain the reference genome pattern ([AT]_x3_[T]_x1_[TA]_x8_), but rather had the [AT]_x3_[T]_x1_[TA]_x7_ variant (**Supplementary Figure S5c-d**). Moreover, this intergenic repeat region was also found to be heteroplasmic in populations that underwent 200 generations of mutation accumulation^36^. Interestingly, a median heteroplasmy percentage of 39% was reported for variant types shifting from [AT]_x3_[T]_x1_[TA]_x6_ to [AT]_x3_[T]_x1_[TA]_x7_ and is relatively close to our median fraction of 29%. In addition to the overlapping finding of the heteroplasmic intergenic repeat region, out of the 32 detected putatively heteroplasmic loci, six more loci overlapped with those reported in the mutation accumulation lines^36^ (**Supplementary Table S16**).

Mitochondrial variants within the DGRP have previously been investigated to study coevolution of the *Wolbachia* endosymbiont and the *Drosophila melanogaster* mitochondria^11^. As part of the validation of our identified mitochondrial variants, we compared our findings. We realigned our data to the compound genome *dm3: chrU, ch3L 10Mb-11.2.Mb* and *Wolbachia* in accordance with previous reports. Using the 134 lines that overlapped between our datasets, we found 115 out of 182 total mitochondrial variants to be present in both datasets (**Figure 2f**). Furthermore, we detected 59 mitochondrial variants that are unique to our dataset. Additionally, eight mitochondrial variants were unique to the previous study^11^. Of these eight, seven were previously considered heterozygous whereas we solely detected the reference allele.

We also assessed whether a correlation exists between the presence of *Wolbachia* and *Drosophila* mitochondrial variants given the coevolution between the two species^11^, but we did not find such relationship (**Supplementary Figure S4**).

### Mitochondrial haplotypes of the DGRP

While the majority of the detected mitochondrial variants were detected in single DGRP lines, 50 variants were detected in 5 or more lines. We therefore postulated that mitochondrial haplotypes may be present within the DGRP. We used the TCS method that employs statistical parsimony and takes the variation within the entire mitochondrial genome into account to separate potential haplotypes^12,29,37,38^. This approach revealed a multitude of haplotypes with most notably, a clear central mitochondrial haplotype, which we dubbed the ‘Central MH’, consisting of 15 DGRP lines. Next, to generate a more refined set of haplotypes, we manually curated the clusters according to the following criterion: a haplotype is formed when a particular variant or set of variants are shared by >5 DGRP lines but not by the Central MitoHap or another haplotype. Lines that could not be placed in distinctive haplotypes were placed in the ‘Outgroup MH’. Using this criterion, we identified 12 haplotypes specific for DGRP lines consisting of an average of 10 lines per haplotype (**Figure 3a; Supplementary Table S17**). The reference strains *w^1118^* and *Oregon-R* each formed their own haplotype.

**Figure 3.**
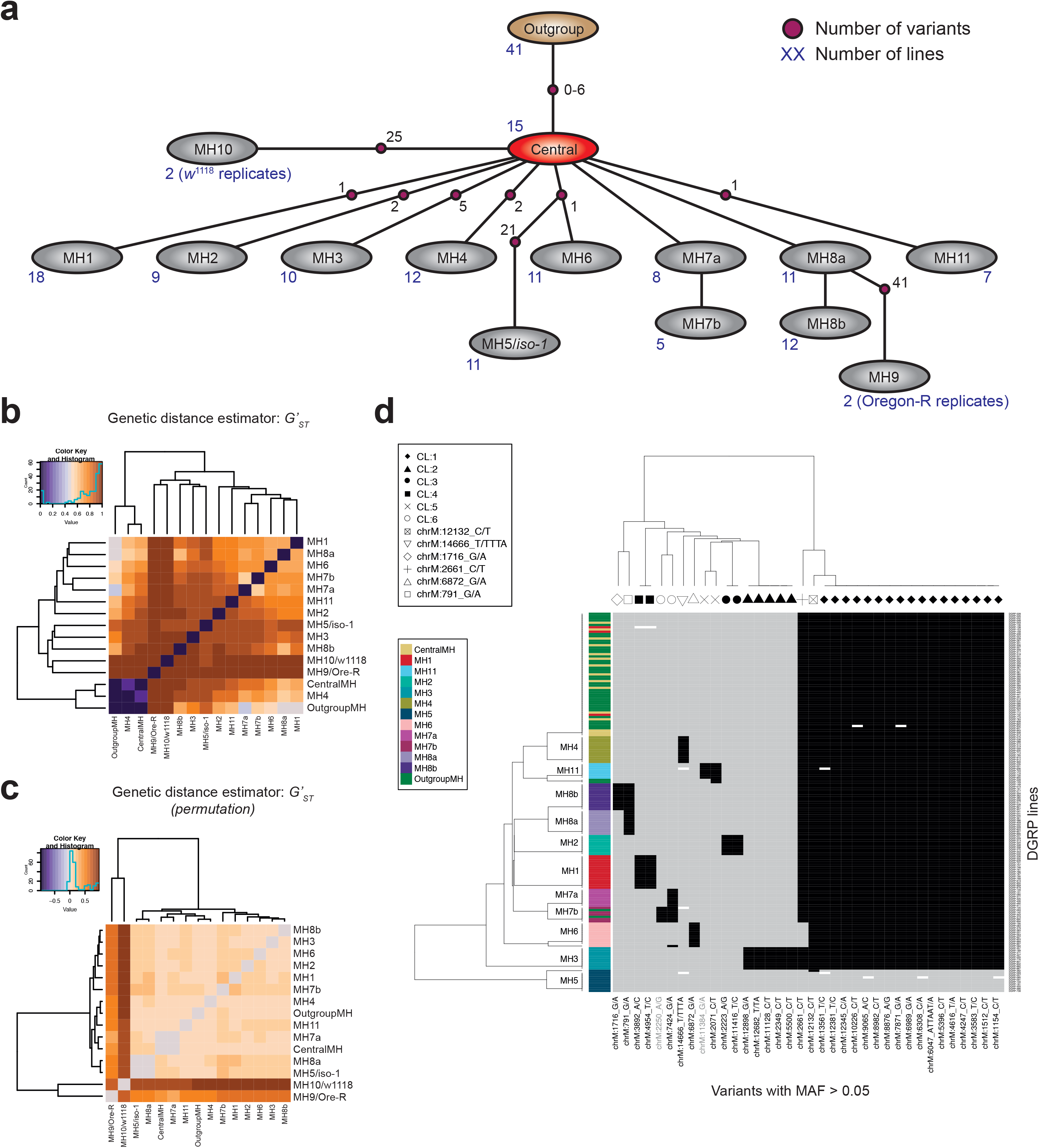
Mitochondrial haplotypes of the *Drosophila* Genetic Reference Panel (DGRP). **a)** Schematic representation of haplogroups based on the TCS method (**Methods**) using multiple sequence alignments. **b)** Measuring the genetic distance between haplogroups from **3a** using the *G’_ST_* estimator. **c)** Permutations of the genetic distance between haplogroups from **3a** using the *G’_ST_* estimator. **d)** Heatmap depicting the haplotype-specific variants. On the bottom x-axis, the variants are depicted with a MAF > 0.05 (black labels) and MAC > 5 (grey labels). On the top x-axis, the cluster-linked variants are indicated with unique symbols. On the right y-axis, DGRP lines are listed with the corresponding haplotypes on the left y-axis. Alternate alleles are depicted with black bars, whereas reference alleles are depicted with grey bars.

We tested the validity of these haplotypes by measuring the genetic distance between them using Hedrick’s *G’_ST_* genetic distance estimator (**Figure 3b**). Overall, the haplotypes clearly separated from one another, implying that the parameters that we used to construct these haplotypes seem to fit. Nonetheless, we also observed three haplotypes MH4, Central MH and the Outgroup MH with a higher genetic relatedness. However, and remarkably, permutation of the DGRP lines of each haplotype resulted in a complete loss of relatedness (**Figure 3c**). These two observations suggest that the TCS method provides a solid basis for the detection of mitochondrial haplotypes in the DGRP.

Furthermore, given that the haplotypes were constructed using the entire set of mitochondrial genomic variants, we assessed whether common variants with a minor allele frequency (MAF) > 0.05 can discriminate the mitochondrial haplotypes. The detection of these common haplotype-specific variants would facilitate further downstream analysis and its usage in phenotype association analyses. Of the 34 variants that have a MAF > 0.05, we detected that approximately half of these are haplotype specific, allowing the separation for each of the detected haplotypes (**Figure 3d**). However, MH7a and MH7b could not be distinguished at the level of MAF > 0.05 because the separating variant had a lower allele frequency (see also **Supplementary Table S8)**. Interestingly, we found that various variants are seemingly linked in particular haplotypes. For instance, we observed five haplotype-specific variants for MH3, and two for MH1, MH2, and MH8b. The relative large physical distance between some of these variants (nearly 10 kb in the most extreme case in MH3) is consistent with the notion of reduced recombination in the mitochondrial genome of *Drosophila melanogaster*^12^.

### Mitochondrial population structure in the DGRP is imprinted on the nuclear genome

The population structure that we observed at the mitochondrial level is particularly interesting given that the DGRP started off from 1,500 isofemale lines obtained from a single geographical location. Many lines did not survive the inbreeding process which may be an indication of selection of either nuclear epistatic effects or mito-nuclear epistatic effects. We therefore investigated whether there is a relationship between mitochondrial and nuclear variants by screening for genotype ratio distortions (GRDs)^30^.

GRDs can be analysed by assessing all mito-nuclear variant allelic combinations which passed the chi-square test on significance (putative GRDs), or in a more conservative manner where a so-called pass-neighbour threshold is applied, meaning that a GRD is only accepted when it has an additional significant GRD down- and upstream within a region of 50 bp. Within the DGRP, the linkage disequilibrium (LD) at this point (50 bp) drops below 0.3^9^. To find large regions affected by GRDs, we first explored the overall landscape of all putative GRDs without applying the pass-neighbour threshold. Due to the linkage between certain mitochondrial variants, these GRDs can largely be considered as mitochondrial haplotype-specific. Therefore, we hypothesized that large genomic haplotype-specific GRD blocks would be detectable if population structure is indeed present.

We indeed observed large regions that were particularly associated with GRDs for a specific mitochondrial haplotype (**Figure 4**). Interestingly, one of the largest regions was located on chromosome 2L from ~2.5 MBp to ~11 MBp containing GRDs associated with MH7. Likewise, on chromosome 3L, we found a region spanning approximately 6 MBp from ~3 MBp to ~9 MBp associated with GRDs for MH1. Moreover, these regions did not seem to be affected by LD, nor by major known inversions that may reside in the DGRP.

**Figure 4.**
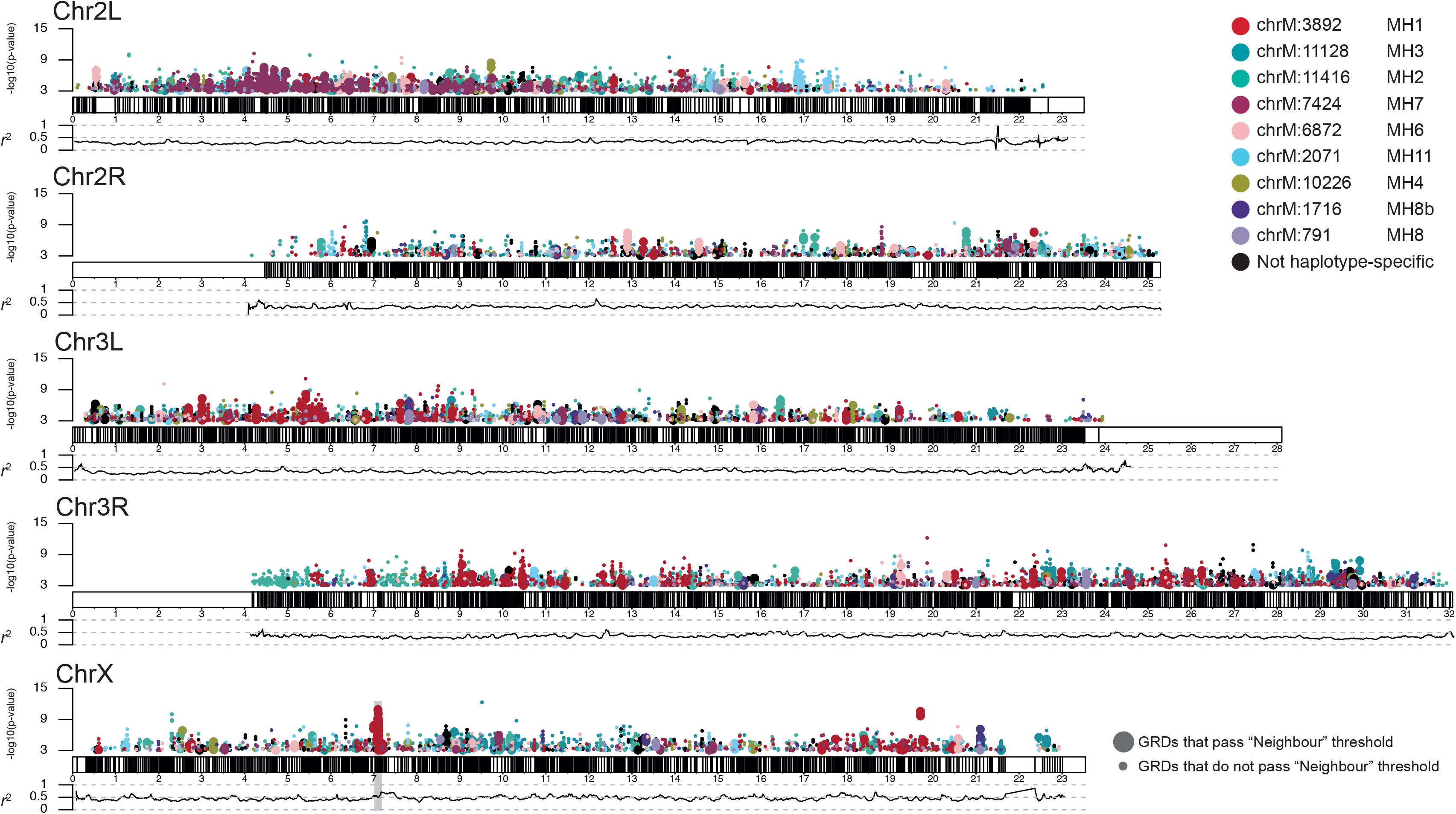
The Genotype Ratio Distortion (GRD) landscape of the DGRP. On top of each chromosome, the location of nuclear variants associated with a GRD is presented. The y-axis presents the −log10(p-value) significance of each GRD. Each colour represents a mitochondrial haplotype. Larger points present those GRDs which pass the neighbour threshold. Below each chromosome, the average LD is presented per mega base in 100kb sliding windows. The *Sex-lethal* locus is highlighted on chromosome X in grey.

Given that we observed large genomic haplotype-specific GRD blocks, we further assessed whether particular haplotypes had more GRDs than others by using GRDs computed with the pass-neighbour threshold. Overall, we detect 1,845 GRDs (mito-nuclear allelic pairs, **Supplementary Table S9**). Most of the GRDs that we detected were associated with MH1 (**Figure 5a**). This is in line with the observation of several large genomic regions associated with GRDs for MH1. While MH1 is the largest haplotype (n = 18 lines), the number of GRDs detected for each haplotype was not correlated with the haplotype size. For instance, MH3 had the second most GRDs (282) whereas the haplotype consisted of 10 lines and MH2 consisting of 9 lines had the third most GRDs (229). When we explored the distribution of significant GRDs over the chromosomes, we found that the greatest density is linked to chromosome 2L (504) and the lowest to chromosome 2R (185) (**Figure 5b, Supplementary Table S10**). This also suggests that the number of GRDs is not affected by chromosome length. Moreover, certain haplotypes seemed specific to particular chromosomes. For example, MH7 (the mitochondrial variant is shared by MH7a and MH7b) was essentially unique for GRDs on chromosome 2L, whereas MH1 was largely spread over chromosomes 3 and X. These findings suggest that the mitochondrial population structure is reflected onto the nuclear genome, and in particular affects specific chromosomes.

**Figure 5.**
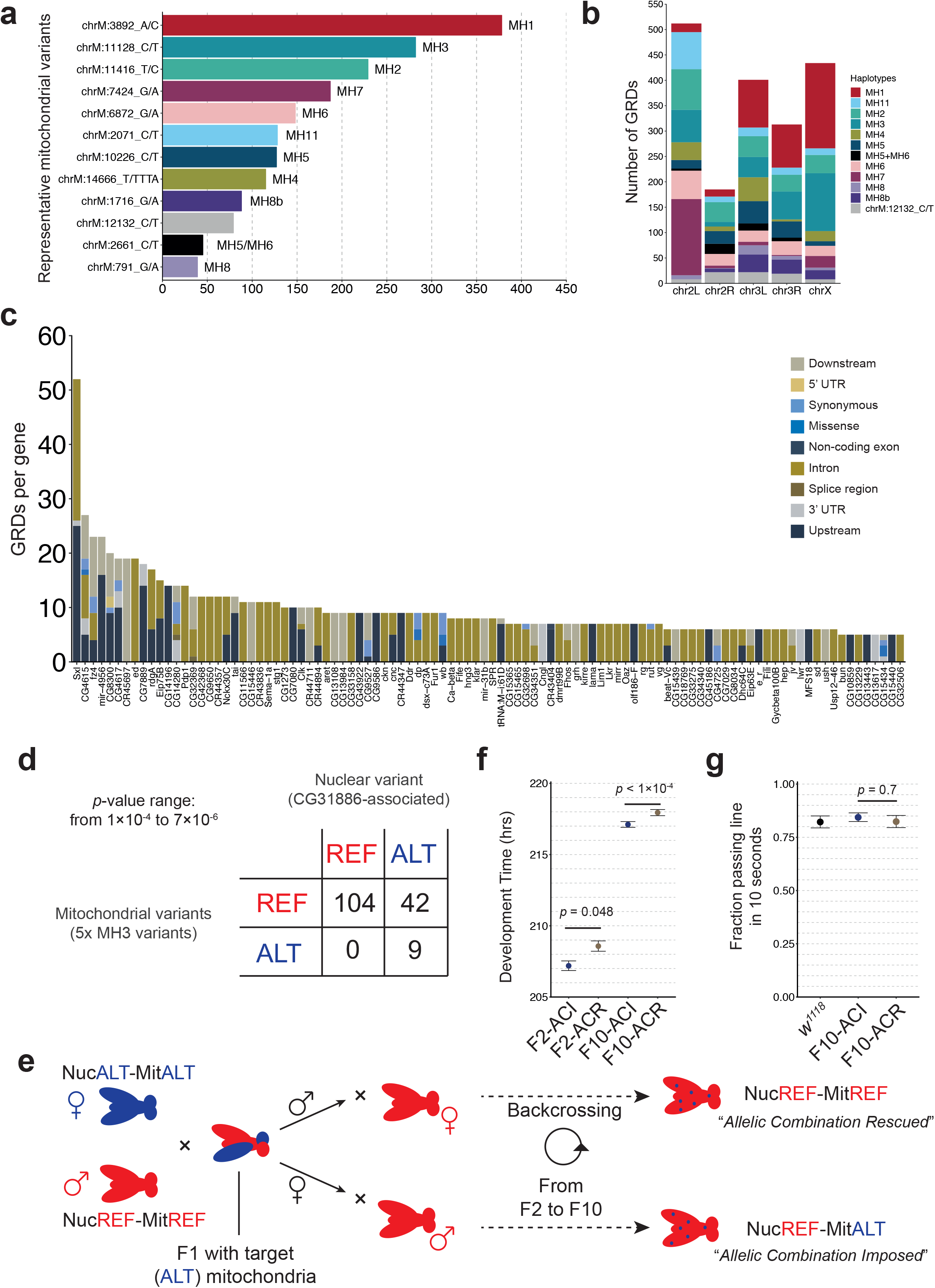
Genotype ratio distortions (GRDs) between mitochondrial and nuclear variants in the *Drosophila* Genetic Reference Panel (DGRP). **a)** Number of GRDs per mitochondrial haplotype. **b)** Distribution of haplotype-specific GRDs per chromosome. **c)** Top 100 genes associated with a GRD. Up/down-stream was set at 2 kb. **d)** Table depicting the variant configuration of the complete GRD (CG31886 perspective). **e)** Schematic overview of the crossing scheme used to produce populations with the absent allelic combination imposed (ACI) or backcrossed and thus rescued (ACR). **f)** Average development time of the F2 and F10 of the imposed (ACI) and rescue (ACR) allelic-combination populations (*p*-values were obtained using Wilcoxon signed-rank tests). See **Supplementary Tables S18-S19** for detailed information on each test. **g)** Climbing activity of imposed (ACI) and rescue (ACR) allelic-combination populations (student’s t-test; t(16) = 0.40, *p* = 0.7).

Given that particular chromosomes were associated with haplotype-specific GRDs, we investigated whether particular genes are proportionally more affected by GRDs and whether genomic regions are reflective of the mitochondrial haplotype. We examined whether particular nuclear genes (and surrounding regions (±2 kb)) are more associated with mito-nuclear GRDs than others. Interestingly, the top gene affected by GRDs was *sex-lethal* (*Sxl*), containing 52 unique mito-nuclear allele pairs (**Figure 5c**). Moreover, we found that the subsequent top 5 genes (*CG4615*, *fz4*, *mir-4956*, *CG8300* and *CG4617*) associated with GRDs are within the same gene region as *Sxl* on chromosome X, indicating that this 40 kb region can be considered as a strong GRD locus (**Supplementary Figure S6**). Interestingly, all of these GRDs were associated with MH1, further supporting our observation that the mitochondrial population structure is reflective of nuclear population structure.

### Genotype ratio distortions do not majorly affect fitness

Most of the GRDs that we detected were so-called ‘incomplete’ GRDs. Incomplete GRDs are those in which all mito-nuclear allelic combinations are present, however the distribution is distorted, whereas in complete GRDs there is a full absence of a particular mito-nuclear allelic combination. To assess whether GRDs have a phenotypic impact, we investigated their effect on fitness phenotypes. In total, we observed 22 complete GRDs, located on chromosomes 2L, 3L, and X (**Supplemental Figure S7**). These were linked to four different haplotype representative variants for MH1, MH3, MH6, and MH11. For MH1, four GRDs were located in the intergenic region between the genes *vvl* and *CR45115*. Six GRDs were detected for intronic variants in the gene *rugose* associated with MH6 whereas three variants in the 5’UTR region of *CG13707* were associated with MH11-specific GRDs. Finally, we detected that MH3 had nine complete GRDs. Five of these were located in the introns of *CG7110*, and one in *rdgA*. While *CG7110*’s function is unknown, *rdgA* encodes a diacylglycerol kinase and has been implicated in odour response^39^, lifespan^40,41^, and starvation resistance^40^. Moreover, and interestingly, the remaining three variants were synonymous SNPs in the gene *CG31886*. Like *CG7110*, the function of this gene is largely unknown, however *CG31886* has been linked to increased ethanol sensitivity^47,48^.

Given that the *CG31886*-GRD associated variants were exonic and that this haplotype displayed the highest number of complete GRDs, we decided to investigate the functional relevance of these complete GRDs. We found that all of the MH3 DGRP lines have the alternate variants for the *CG31886* locus, and thus have the same allelic configuration (**Figure 5d**). We hypothesized that if an allelic combination of a locus is truly genetically incompatible, we would expect a decreased fitness. We assessed if DGRP lines would show developmental or activity defects. Briefly, we crossed females of DGRP lines containing the mitochondria of interest with males that had the nuclear reference background (**Figure 5e**). This resulted in the generation of F1 lines that had the ‘alternate’ mitochondria with a heterozygous nuclear background. The F1 females were used to build populations in which the allelic combination of interest was imposed (‘AC-imposed’ = ACI) by backcrossing with males containing the nuclear reference background. The F1 males were used to backcross populations to their original nuclear background (‘AC-rescue’ = ACR). These AC-rescue lines would thus retain residual nuclear fragments of the ‘alternate mitochondrial’ lines making the comparison fairer rather than using the parental DGRP lines directly. Further backcrossing while maintaining the mitochondria of interest (or rescue) resulted in F10 populations (see **material and methods**, and **Supplementary Figure S2** for a more detailed description). This resulted in the generation of nine lines (3×3 parental strains) that had the allelic combination imposed (‘AC-imposed’) and nine lines that were backcrossed towards their original state (‘AC-rescue’). Any phenotypic variation between an imposed and a rescue line should theoretically be the result of a specific mito-nuclear genetic interaction.

To assess overall fitness, we measured the development time of the F2 and F10 lines. In the F2 generation, we obtained the first generation of flies that have a homozygous nuclear background for the locus of interest. However, since the nuclear background of the F2 flies is still largely mosaic, we also further backcrossed to F10. Contrary to expectations, we observed a small but significant effect in the development time where the imposed-AC crosses developed faster than the rescue-AC group in the F2 generation (Wilcoxon; *p* = 0.048) and a large significant effect in the F10 generation (Wilcoxon; *p* < 1×10^−4^) (**Figure 5f**). Furthermore, we did not observe clear differences in sex ratio or eclosion (**Supplementary Figure S8a-g** and **Supplementary Table S18-S20**), nor did we observe a significant effect in the climbing activity of the F10 generation flies (**Figure 5g** and **Supplementary Figure S9**).

The lack of clearly negative fitness effects linked to this complete GRD may be an aberration, which is why we aimed to more systematically assess the phenotypic impact of GRDs. To do so, we systematically screened GRDs for any association with previously published fitness traits (see **Methods** section). Surprisingly, we did not observe any strong phenotypic associations, since out of the 29 fitness phenotype groups that were tested among eight groups (haplotype enrichment in 4x mito-nuclear allelic combinations and 2x top and bottom groups) in 1,845 GRDs, we only detected three significant hits. In these cases, paraquat resistance in females^42^ was associated with GRDs from the alternate allele of chrM:7424_G/A (MH7) and three nuclear reference alleles (2/3 were in LD). The nuclear variants were all intronic and, interestingly, the genes associated to them were *rutabaga* and *REPTOR*. The latter is of interest since knockdown of REPTOR, a repressor of the TORC1 complex, has been linked with decreased starvation resistance in *Drosophila*^43^.

### Integration of mitochondrial haplotypes in phenotype association analyses

Whereas GRDs appeared to have limited functional impact, individual mitochondrial variants may still contribute to phenotypic variation. To formally address this possibility, we investigated the role and impact of mitochondrial variation and population structure in genome-wide association (GWA) analyses for a compendium of >600 publicly accessible DGRP phenotypes that we assembled for this purpose, and that we reduced to 259 non-correlated, independent phenotype groups (**Methods**). Out of 231 mitochondrial variants, we only considered 34 since the latter passed the threshold of a MAF >0.05 based on the total number of lines sequenced, often resulting in 8 lines, adding to the statistical power (**Figure 3d**). Since we showed that these variants effectively reflect the haplotypes found via the TCS method, we decided to simplify the envisioned analyses and to explore associations between the 259 stipulated phenotype groups and the mitochondrial haplotypes as explanatory variables. We applied a two-stage procedure to detect differences in phenotypic means between haplotypes. First, we applied a relatively lenient ANOVA test to each phenotype group to filter phenotypes that are potentially influenced by mitochondrial haplotypes. Second, if the focal phenotype passed our initial screen, then we applied the TukeyHSD test to identify the significant pairwise differences between phenotype means of haplotypes after which the p-values were adjusted for multiple testing. Using this approach, we identified 12 phenotypes with a significant difference between at least one pair of haplotypes (**Figure 6a** and **Supplementary Table S13**). Most notably, metabolism-related phenotypes such as food intake in males^34^ (MH1 vs MH5/iso-1, *p* = 0.04), waking activity CVE^44^ (MH8b vs MH3, *p* = 0.02), and amount of triglycerides on high glucose diet^45^ (MH2 vs MH7a, *p* = 0.04) were among the top significant hits.

**Figure 6.**
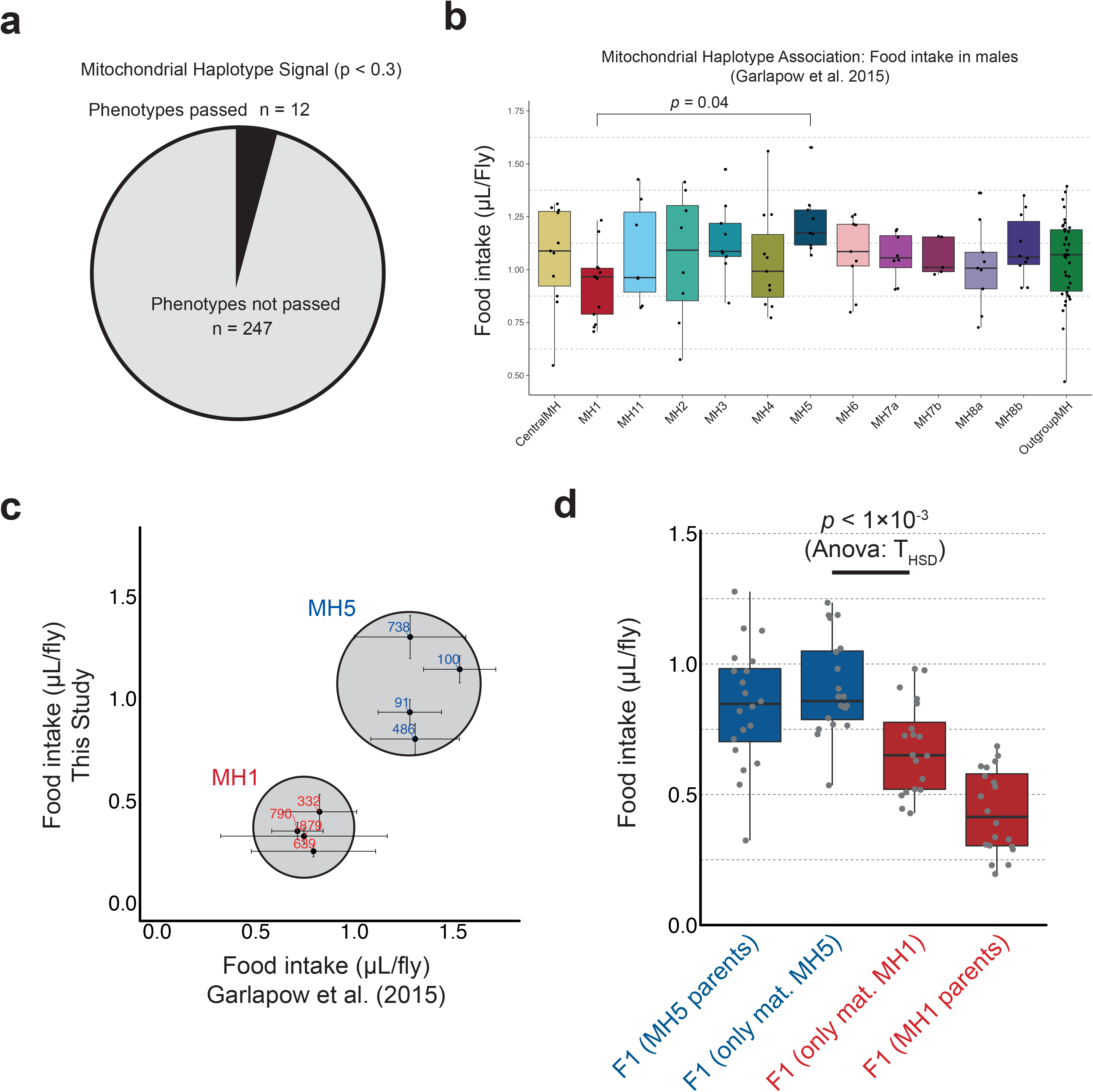
Mitochondrial haplotype association (MHA) analysis. **a)** Fraction of phenotypes passing the ANOVA screening stage of the MHA analysis. **b)** Food intake in males (Garlapow *et al*. (2015)^34^)showing a significant difference between haplotypes MH1 and MH5. **c)** Reproducibility of the results from Garlapow et al. (2015). On the y-axis, the food intake as measured in this study and on the x-axis, the food intake as reported by Garlapow et al. (2015). In blue, the food intake is shown for DGRP lines from haplotype MH5 (high food intake) and in red from haplotype MH1 (low food intake). **d)** Box plots showing the food intake for crosses that either had MH1 (red) or MH5 (blue) mitochondria.

To experimentally validate the findings of the haplotype-based association analysis, we focused on food intake in males given that the two haplotypes (MH1 and MH5/iso-1) were also at the extreme ends of the phenotype distribution (**Figure 6b**). Interestingly, we observed that the effect size of the MH1-MH5/iso-1 mitochondrial haplotypes (1.76, CI = 0.7-2.81) is higher than most of the detected nuclear variants in the reference study by Garlapow et al. (2015)^34^ and lies in the 97th quantile of the absolute effect size distribution of nuclear variants (min = 1.095, max = 2.325, median = 1.275)^34^. This provides further support to our postulate that the mitochondrial genome influences food intake. It must be noted, however, that the effect size calculations could be misrepresented given that we were able to employ 13 groups (haplotypes) versus only 2 groups (REF vs. ALT allele) in the association analyses.

First, we assessed whether we could reproduce the food intake measurements reported by Garlapow et al. (2015)^34^ for selected lines from the phenotypic extremes (**Figure 6c**). In general, our results largely recapitulated those of the reference study^34^. To test whether we could increase the food intake of low phenotype lines by swapping their mitochondrial genomes with those of high food intake lines, we selected two high food intake lines and two low food intake ones which we crossed with between haplotypes and within haplotypes. We measured the food intake of the F1 and found that the presence of mitochondria linked to high food intake indeed induced higher food intake in otherwise low food intake lines and that the pooled effect of the crosses was significant (*p* < 1×10^−3^; ANOVA and THSD; **Figure 6d**). Furthermore, we found that the effect size for the mitochondrial haplotypes is 1.3 (Cohen’s *d*) and thus marginally larger than the median effect size of the nuclear variants. Since this assay was performed in males, it is possible that there is an effect from the X chromosome from either of the haplotype lines, although no significant X-linked variants were associated with high or low food intake by Garlapow et al. (2015)^34^. In sum, when considering the raw difference in food intake between the F1 crosses, then we observed an average difference in food intake of 0.234 µL per fly as a result of mitochondrial haplotypes. Thus, using food intake in males as an example phenotype, these results demonstrate the value of integrating mitochondrial haplotypes in association studies to inform on the genetic determinants of quantitative traits.

## Discussion

The *Drosophila* Genetic Reference Panel (DGRP) has been used to identify nuclear genetic determinants for many traits, but the effect of mitochondrial variation has remained largely unexplored. Here, we expanded on current knowledge of mitochondrial variation in the DGRP by sequencing the mitochondrial genomes of 169 DGRP lines using a robust and efficient mtDNA-enriched sequencing method. With this method, we were able to generate a high-resolution catalogue of mitochondrial genomic variants. In comparison to Richardson *et al*. 2012^11^, we detected 51% more mitochondrial variants, totalling 231 variants in the coding regions of the mitochondrial genome of the DGRP. It is unlikely that any of these new variants occurred *de novo* in our lab and became fixed, given that multiple newly detected variants are present in higher allele frequencies. Genes that were more affected by missense than synonymous variants were *mt:ATPase6* and *mt:Cyt-b*. This is in line with earlier reports for *mt:ATPase6*^46^ and *mt:Cyt-b*^47^ in which it was shown that in specific *Drosophila melanogaster* populations, though not all^48^, both genes are more prone to amino-acid substitutions than in other species, reflecting perhaps a greater tolerance to genetic variation. Finally, we resolved a particularly challenging putatively heteroplasmic intergenic repeat region that Haag-Liautard *et al*. 2008^36^ also reported to be heteroplasmic at levels of up to 39% in mutation accumulation populations. In our study, we found this region to be heteroplasmic in 120 DGRP lines at a median level of 28% of the reads containing an alternate allele. The findings of Haag-Liautard *et al*. 2008^3^6 and our study suggest that heteroplasmy at the population level in *Drosophila* is common, and that particular loci are more prone to be heteroplasmic than others.

We resolved 12 mitochondrial haplotypes for DGRP which we validated based on their genetic relatedness and permutation analyses. Nevertheless, we also observed that DGRP lines containing particular haplotype defining mitochondrial variants were assigned to the Outgroup MH. This can be attributed to the fact that the TCS method utilizes the entire mitochondrial genome rather than estimating the relatedness solely on individual variants, and thus other variants may have been the cause of this. Furthermore, we observed that certain mitochondrial variants were linked which is in line with observations that recombination in the mitochondrial genome is low^12^. While the haplotypes are indicative of population structure at the mitochondrial level, this would further suggest that this population structure was already present at the time when the isofemale lines were established.

The nuclear and mitochondrial genome need to cooperate for a cell or organism to function properly^2,3,49,50^. Given that we detected strong mitochondrial population structure, we investigated the presence of genotype ratio distortions (GRDs) between nuclear and mitochondrial alleles. Overall, we found 1,845 significant GRDs, a number that is far greater than that reported by Corbett-Detig et al. 2013^30^. However, their study focused on inter-chromosomal autosomal GRDs. Interestingly, we found that these GRDs disproportionally favour particular mitochondrial haplotypes and that these haplotypes in turn favour particular chromosomes. Moreover, many of the GRDs tend to form ‘blocks’ of multiple nuclear variants along a sequence, indicating that the nuclear genome seems to be imprinted by the population structure that is observed at the mitochondrial level. This can for instance be observed from the large genomic haplotype-specific GRD blocks in the GRD landscape (some several megabases long) and also notably at a smaller scale in the *Sxl*-GRD locus associated with haplotype MH1 and the *CG31886/rdgA*-GRD with MH3).

Given the large number of GRDs, we also explored which genes or genomic regions are mostly affected. We initially revealed a 40 kb region around the gene *sex-lethal* (*Sxl*) to be primarily linked with GRDs. Alterations in *Sxl* can be pivotal to sex determination and can be lethal to females^51^. Furthermore, Tower (2015)^52^ postulated that *Sxl* is implicated in mitochondrial maintenance via dosage compensation, suggesting that specific combinations of mitochondrial and *Sxl-*linked variants could modulate mitochondrial homeostasis. Additionally, of the approximately 1,900 nuclear encoded genes linked to a GRD, 120 were mitochondrial-associated (6.3%). This is roughly the same proportion as the estimated mitochondrial-associated genes in the genome over the total number of genes (7.5%). We therefore speculate that GRDs do not disproportionally affect nuclear encoded mitochondrial genes.

We assessed the functional relevance of these GRDs by experimentally analysing the effect of one complete GRD, associated with MH3 and related to the gene *CG3886*, on development time and climbing activity. Contrary to expectations, we observed a significant decrease in development time and no significant difference in climbing behaviour commonly associated with mitochondrial function. These results suggest that the imposed “unnatural” allelic combinations do not induce functional deficits. We cannot exclude at this point that the utilized laboratory conditions may have masked certain effects or that other, unexamined phenotypes could be affected. This is why we performed an additional systematic phenotype-GRD correlation screen. However, despite the high number of detected GRDs, we could again not find any strong signal besides the three associations with paraquat resistance in females. One other possibility for the lack of strong phenotypic effects is that these GRDs are false positives. This is unlikely though because our analysis was conservative in nature and both the mitochondrial and nuclear variants used here were filtered out if the allele frequency was too low (MAF > 0.05). Thus, if the GRDs are accurate, then there may be other reasons for this.

Specifically, we postulate that these GRDs appear in our analysis as the result of an initial non-lethal population structure, where a subset of isogenic females at the start of the collection had this allelic combination of mitochondrial and nuclear variants. In other words, the detected GRDs may be more likely the result of standing population structure when establishing the DGRP rather than representing actual genomic incompatibilities. Indeed, the DGRP started off from a collection of more than 1,500 isofemale lines, but through the process of full-sibling mating, only 200 remained viable and could be maintained^53^. The current DGRP consists therefore mainly of robust fly lines that are not affected by highly deleterious, fitness-reducing loci, which in turn may rationalize why we do not observe large GRD-linked effects from the GRD-landscape. However, our results may also indicate that mito-nuclear GRDs have in general less influence on phenotypic variation compared to nuclear genomic GRDs, which have already been shown to affect fitness traits such as development in *Drosophila*^54^. This rationale would be consistent with recent findings reported by Mossman et al. (2016), who showed that DGRP lines subjected to introgression of various mitochondrial genomes^15^ exhibit development phenotypes across certain diets, but overall remain viable, to the extent that even introgression of mitochondria of the sister species *Drosophila simulans* are tolerated.

This raises the question to which extent individual mitochondrial haplotypes actually impact on phenotypic variation. While it has been shown that such haplotypes can affect life history traits^14^, a systematic analysis investigating the extent of mitochondrial variation which underlies phenotypic variation has so far been lacking, especially using controlled populations with a large number of phenotypes. To address this, we performed a mitochondrial haplotype association analysis involving 259 uncorrelated phenotypes, which revealed 12 that contained haplotype pairs that significantly differed from one another. Interestingly, a substantial proportion of these phenotypes are linked to metabolism or stress response, which is intuitive because of the well-appreciated role that mitochondria have in these biological processes^55,56^. As a proof-of-concept, we validated these results on food intake in males, demonstrating to our knowledge for the first time that mitochondrial haplotypes can affect feeding behaviour, although in a sexually dimorphic fashion.

## Acknowledgements

We would like to thank the lab members of the Deplancke lab for helpful suggestions on experiments and analyses. Especially Dr. Daniel Alpern for help with the Tn5 protocol, Dr. Vincent Gardeux and Michael Beyeler for help with the sequence and phenotype association analysis, Michael Frochaux for experimental assistance and Dr. Antonio Meireles-Filho for rendering the Tn5 transposase available. We further thank Dr. Bastien Mangeat (EPFL, GECF) and Dr. Keith Harshman (UNIL, CIG) for the sequencing of the libraries, and Dr. Bianca Habermann for valuable suggestions on the mito-nuclear genes. We are also very grateful for the computational infrastructure provided by the Swiss Institute of Bioinformatics and by Vital-IT at EPFL. This project was funded by a grant from SystemsX.ch to BD (AgingX) and by Institutional Support of the EPFL.

## Data availability

The sequencing data is available at NCBI Sequence Read Archive (SRA) is currently submitted under SRP168326 (see also **Supplementary Table S2**). All other data (i.e. Variant files, GRDs) can be found in the remaining Supplementary Tables.

## Author contributions

Conceptualized the study RPJB, ML, and BD. Performed the experiments: RPJB, ML, and VSB. Performed the computational analyses ML, RPJB, and AK. Critical suggestions and comments on manuscript: MRR, JA and BH. Wrote the manuscript RPJB, ML, and BD.

---

**Supplementary figure S1.**
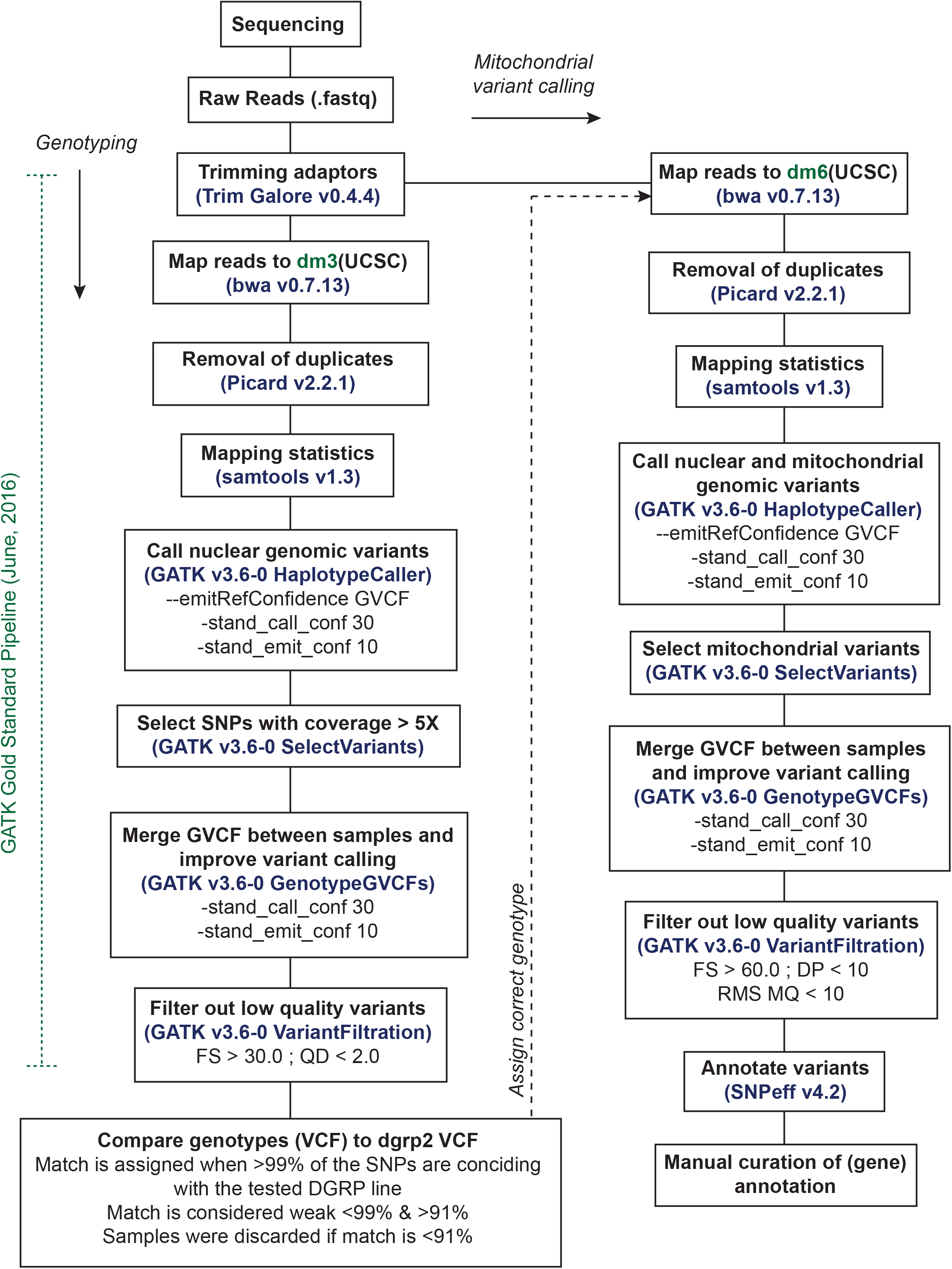
Flow scheme of genotyping and variant calling. Samples were first genotyped based on nuclear fragments which were sequenced. Corrections for the genotype (where necessary) were applied prior to mitochondrial variant calling.

**Supplementary figure S2.**
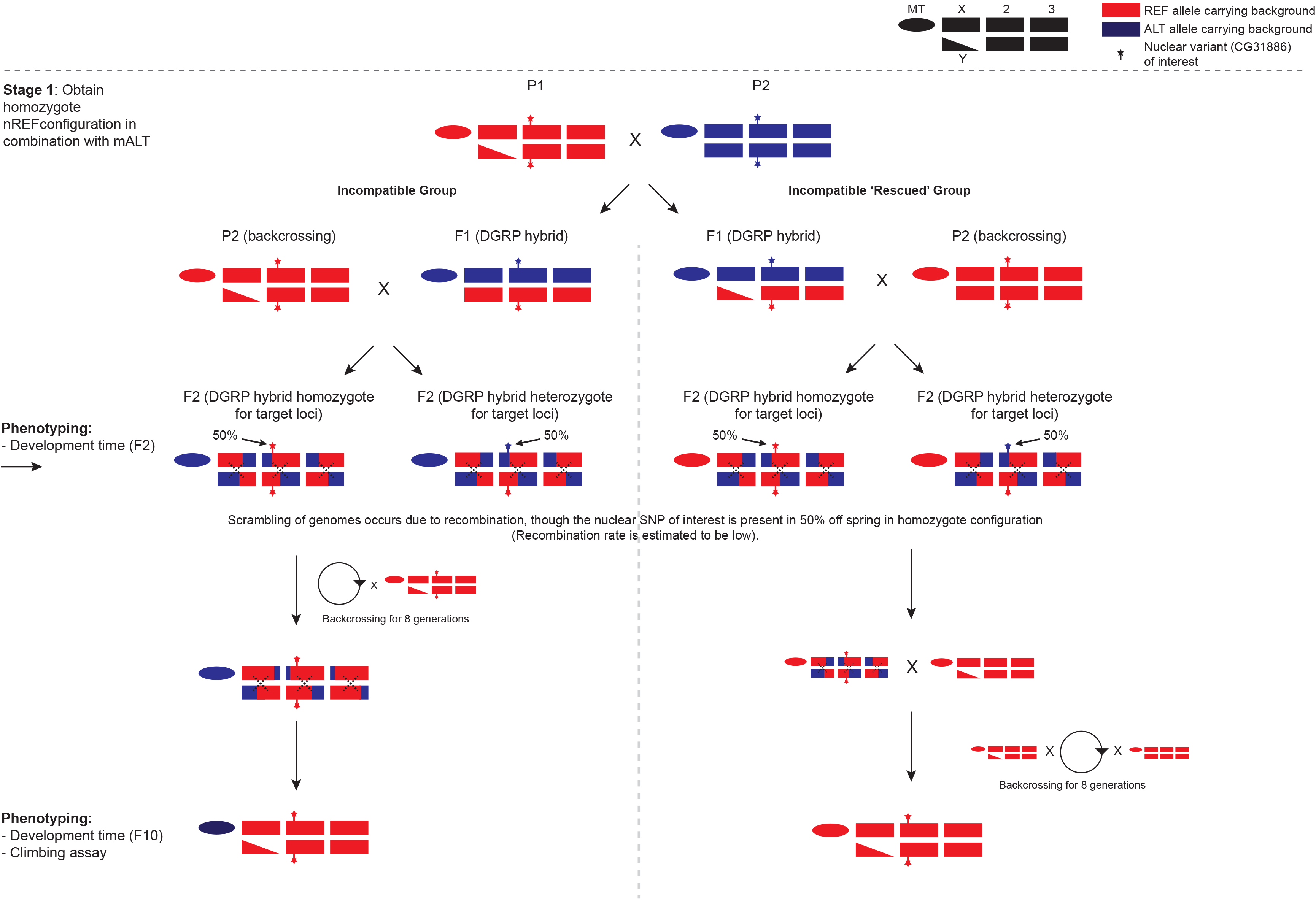
Crossing scheme to study mito-nuclear incompatibilities. Each block represents an autosome and ovals represent the mitochondrial genome. The most left block represents the sex chromosomes, where males are depicted with a triangle and a block, and females are depicted with two blocks (see also upper right legend). Lines with the reference allele (REF; in red) in the mitochondrial and nuclear genome were crossed against lines with the alternate allele (ALT; in blue) in the mitochondrial and nuclear genome. The nuclear variant of interest is depicted with a star on the nuclear genome. The target configuration is ALT mitochondrial genomes in a REF nuclear genomic background (Incompatible group). After the F1 generation, offspring were split where one group (females) was further backcrossed with the REF nuclear background to generate REF homozygous backgrounds. The remaining group (males) were backcrossed for 2 generations with REF mitochondrial and nuclear genomic females to reintroduce the REF mitochondrial genome, before backcrossing with REF mitochondrial and nuclear genomic males was continued. Development assays were performed at generation F2 and F10, and the climbing assay only for F10.

**Supplementary figure S3.**
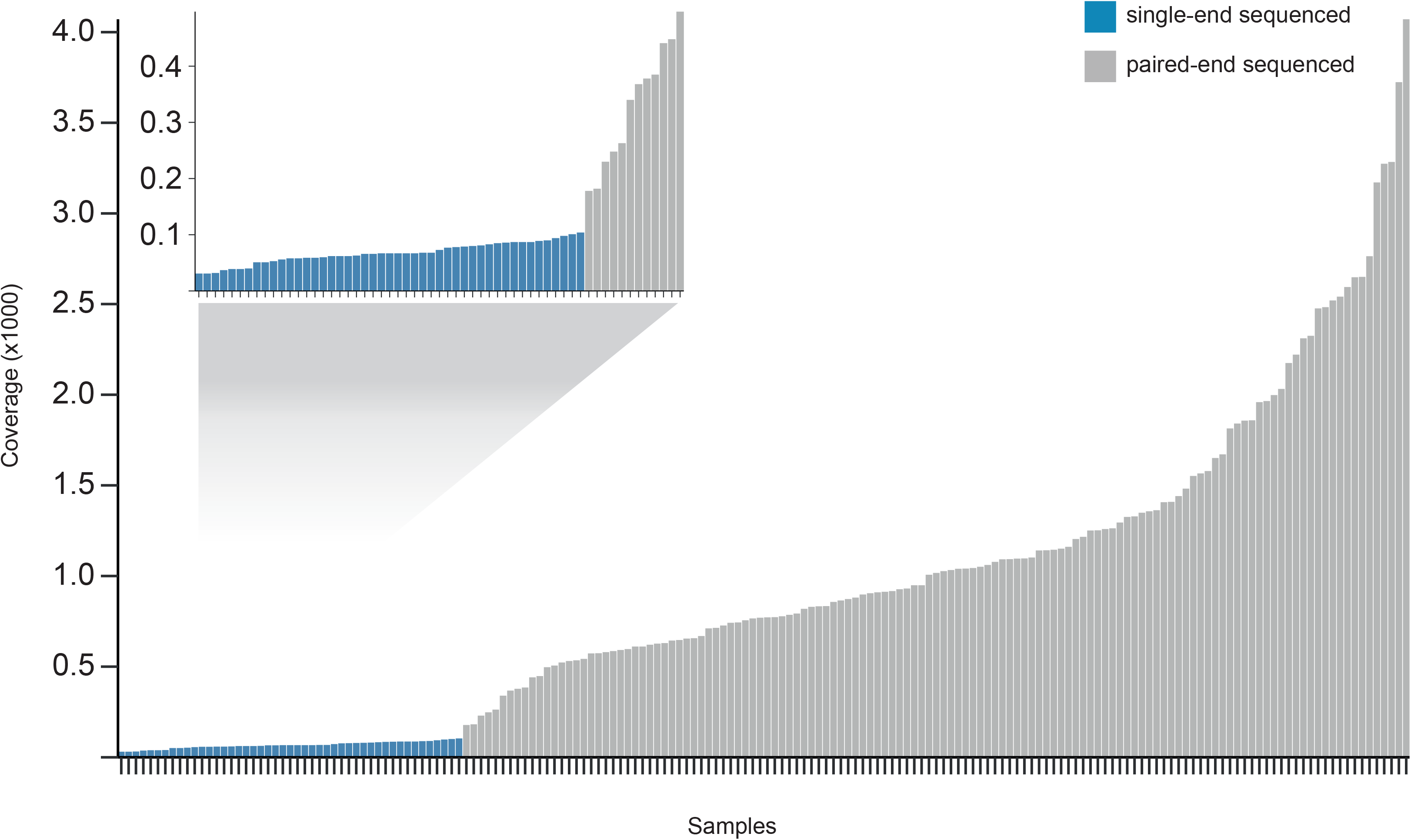
Coverage of mitochondrial DNA per sample. Each bar represents the coverage of a single DGRP line. Samples sequenced using a paired-end strategy are shown in grey and samples sequenced using a single-end strategy are shown in blue. See also **Supplementary Table S4** for a detailed overview of sequence statistics per sample.

**Supplementary figure S4.**
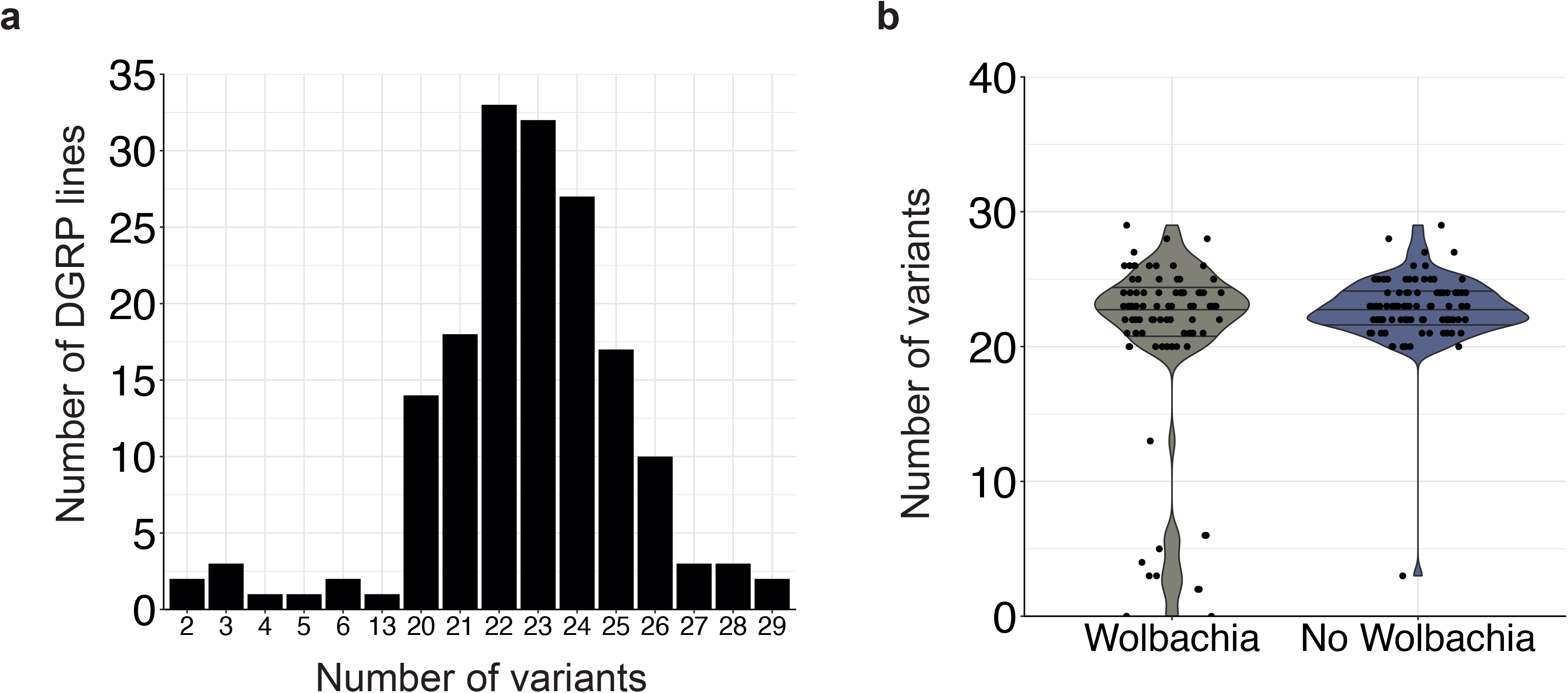
Average number of variants and the relationship to *Wolbachia* infection. **a)** Average number of mitochondrial variants per DGRP line. **b)** Relationship between the number of mitochondrial variants in DGRP lines and *Wolbachia* infection.

**Supplementary figure S5.**
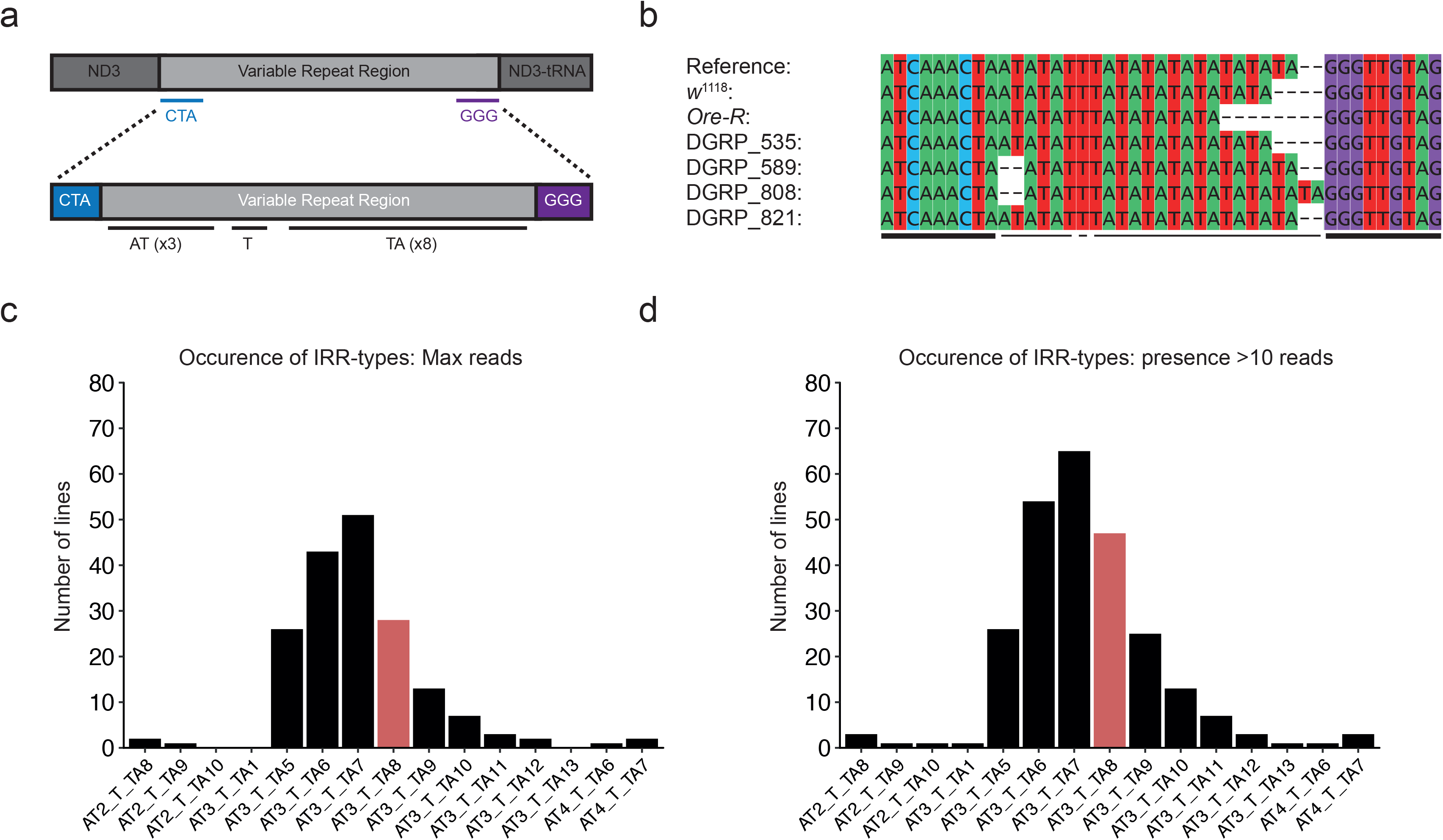
Mapping of a putatively heteroplasmic intergenic repeat region in mitochondrial genomes of *Drosophila* Genetic Reference Panel (DGRP) lines. **a)** Flanking regions that were used to accurately retrieve reads containing the heteroplasmic repeat sequence. **b)** Sanger sequencing of randomly selected DGRP lines of the intergenic repeat region showing the diversity between lines. **c)** Frequency of the dominant (maximum number of reads) types of the heteroplasmic intergenic repeat region in the DGRP. In red is the reference type (carried by *iso-1*). **d)** Frequency of all types of heteroplasmic intergenic repeat region observed in all DGRP lines that are supported by 10 or more reads. In red is the reference type (carried by *iso-1*).

**Supplementary figure S6.**
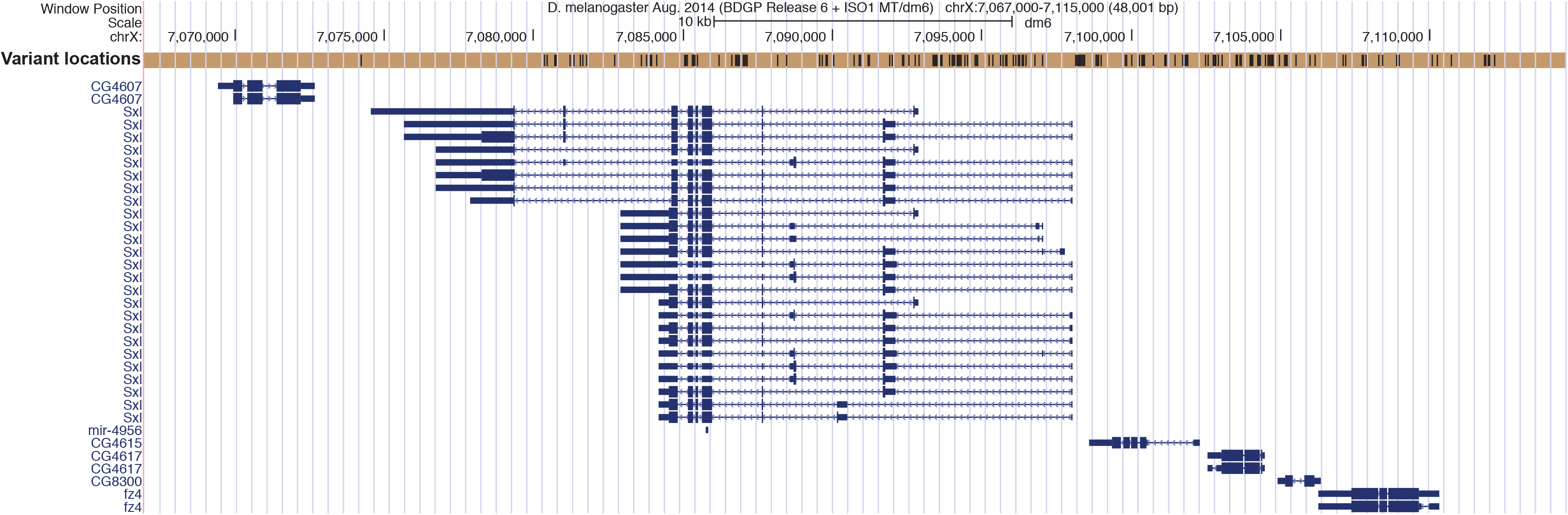
Gene region of Sex-lethal (Sxl) and surrounding genes. Locations of nuclear genomic variants in the DGRP are shown above the genes.

**Supplementary figure S7.**
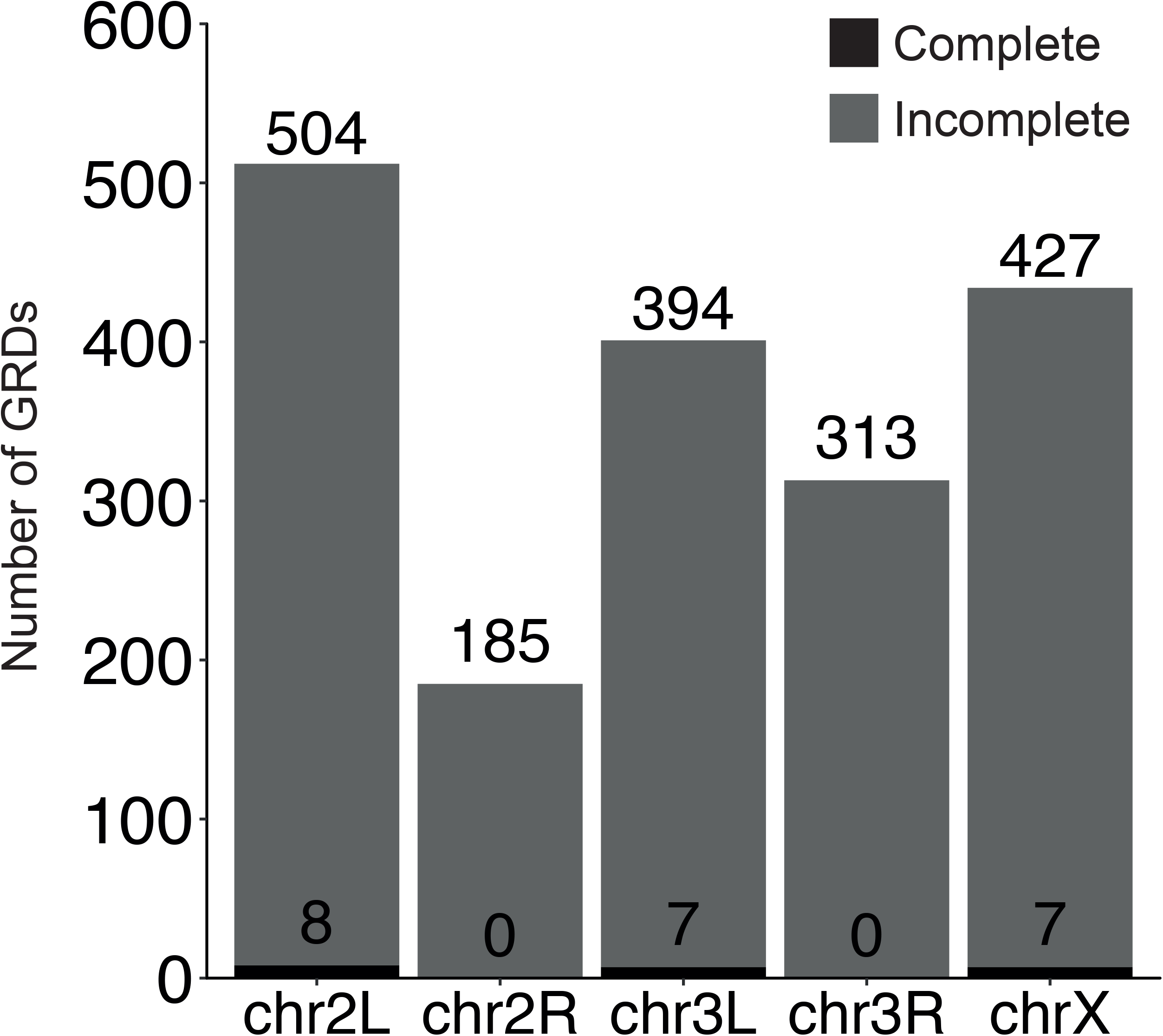
Number of complete and incomplete Genotype Ratio Distortions (GRDs).

**Supplementary figure S8.**
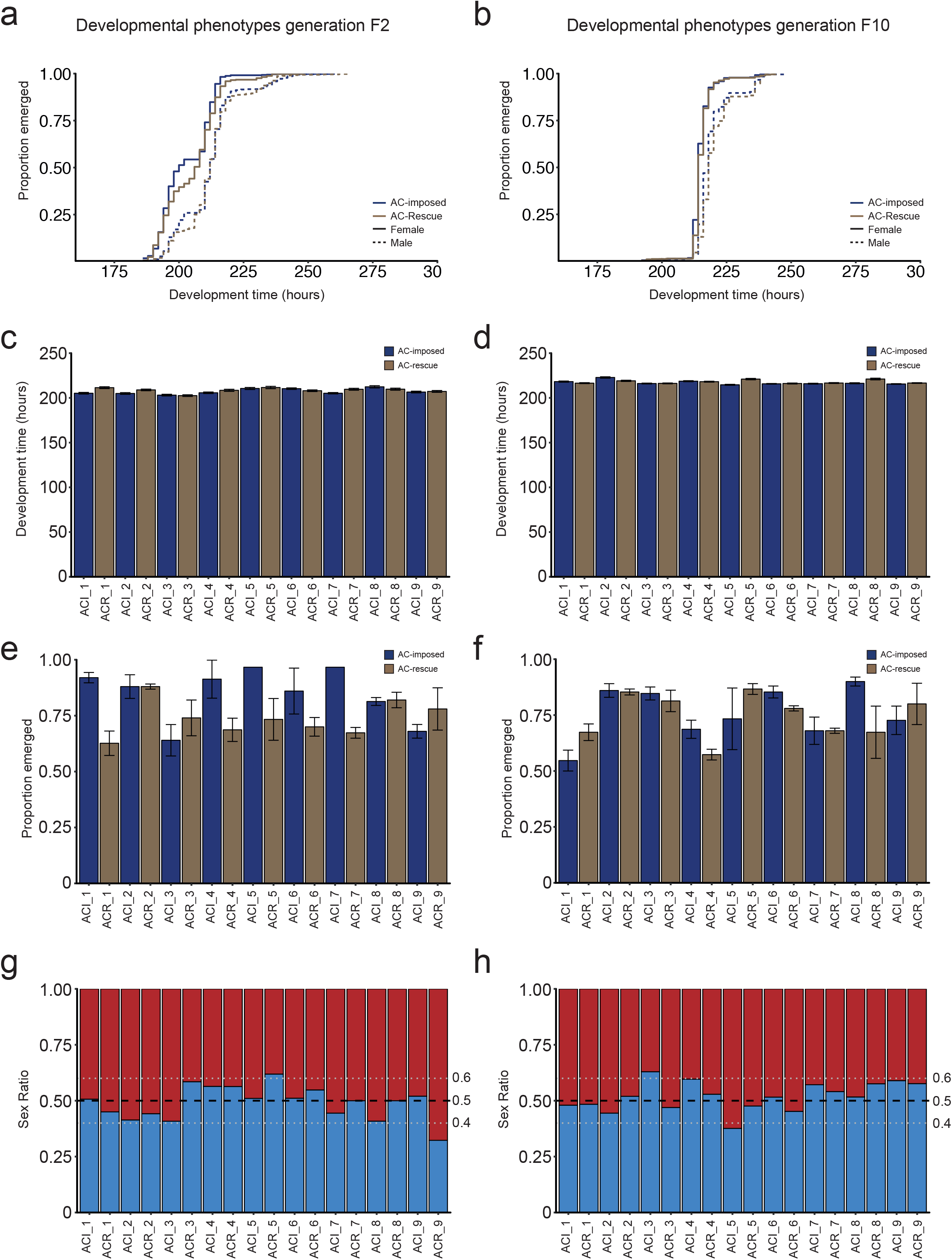
Secondary developmental phenotypes of F2 and F10 imposed allelic combination populations. **a,c,e,g)** Results for the F2 generation of the imposed and rescue allelic populations. **b,d,f,h)** Results for the F10 generation of the imposed and rescue allelic populations. **a-b)** Development time curves. In purple, the imposed AC populations are presented and gold the rescue AC populations. Solid lines represent the development curve for females and dashed for males. **c-d)** Average development time for individual populations. **e-f)** Proportion of flies emerged for individual populations. **g-h**) Sex-ratio of the emerged flies per population.

**Supplementary figure S9.**
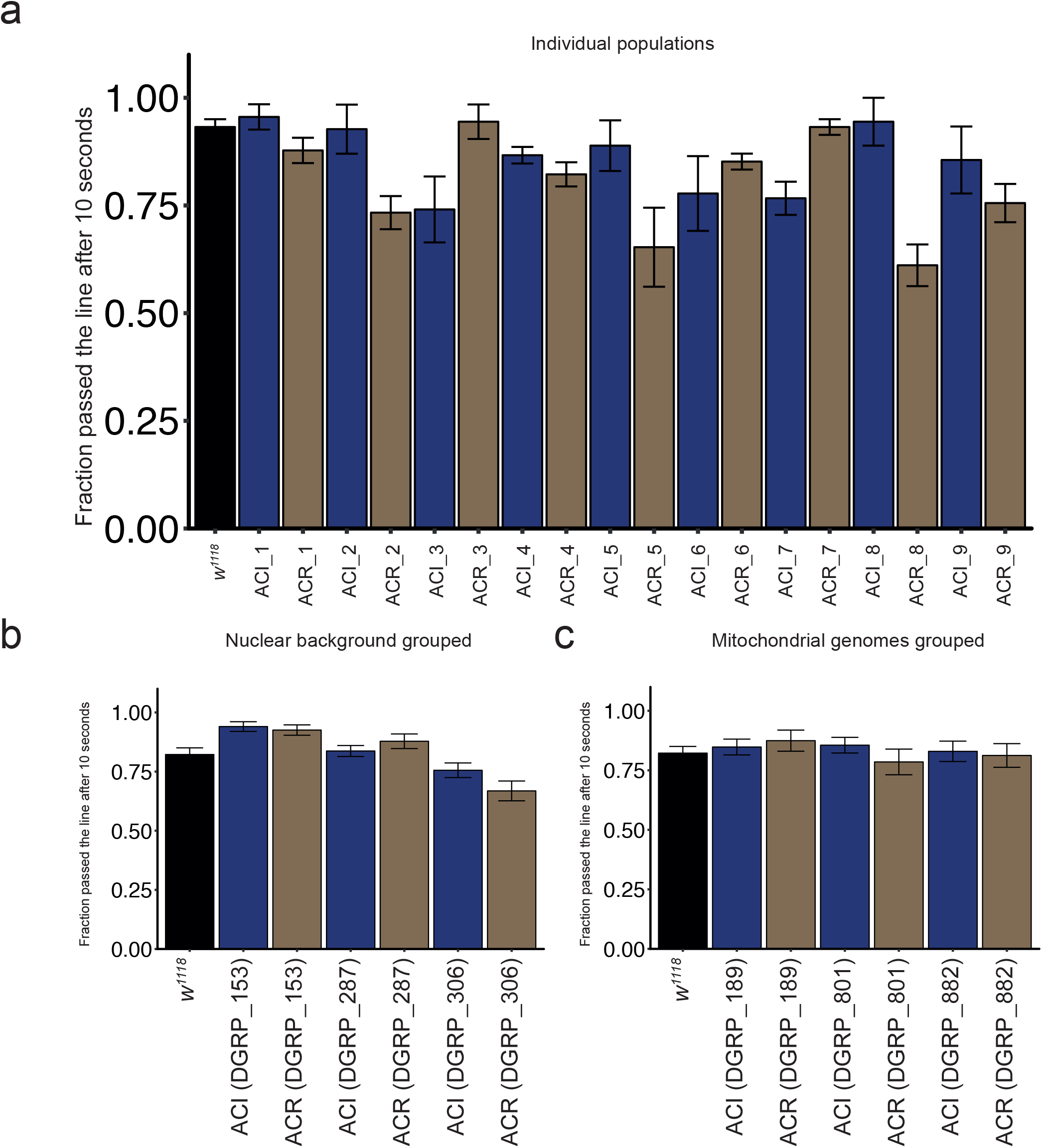
Climbing capability of imposed and rescue allelic combination populations. **a)** Fraction of flies that passed the line after 10 seconds for each of the tested populations. **b)** Fraction passed from the perspective of nuclear backgrounds and **c)** mitochondrial background.

**Supplementary table S1. List of DGRP lines used in our study.**

**Supplementary table S2. Accession numbers of sequencing data.**

**Supplementary table S3. List of primers used in our study.**

**Supplementary table S4. Sequence statistics.**

**Supplementary table S5. Sanger sequencing primers.**

**Supplementary table S6. Sanger sequencing results.**

**Supplementary table S7. Mitochondrial variant table.** 0 = reference, 1 = homozygote alternate, 2 = heterozygote alternate, 3 = homozygote for the second alternate 4 = heterozygote for the second alternate.

**Supplementary table S8. Mitochondrial variant clusters and allele information.**

**Supplementary table S9. All mito-nuclear Genotype Ratio Distortions in the DGRP.**

**Supplementary table S10. Summary of Genotype Ratio Distortions.**

**Supplementary Table S11. Lines used for the CG31886-rdgA complete GRD crosses.**

**Supplementary table S12. Source of fitness phenotypes used.**

**Supplementary table S13. Mitochondrial Haplotype Association top hits.**

**Supplementary table S14. Number of lines that have (multiple) nucleotides with a coverage below threshold.**

**Supplementary table S15. Total composition of mitochondrial variants within the DGRP.**

**Supplementary table S16. Overlap of heteroplasmic loci with Haag-Liautard et al. 2008^36^.** [a] chrM:IRR refers to the intergenic repeat region in which we detect variation in the variant types and levels of heterozygosity. [b] Our variant caller finds three versions and hence only first T, TA (TA1), and TAA (=TA2) are considered as variants rather than TA8 or TA9.

**Supplementary table S17. Mitochondrial haplotypes of the DGRP.**

**Supplementary table S18. Log-Rank and Wilcoxon survival analysis statistics on the development time.** * < 0.05, ** < 0.01, *** < 0.001, **** < 0.0001.

**Supplementary table S19. Average development time per population group.**

**Supplementary table S20. Average development time for each individual subpopulation.**

